# The actin-associated protein Kaptin modulates F-actin barbed-end dynamics

**DOI:** 10.1101/2023.10.30.564586

**Authors:** Priyanka Dutta, Ipshita Maiti, Aurnab Ghose, Radha Chauhan, Sankar Maiti

## Abstract

Living cells require a dynamic and precisely regulated actin cytoskeleton to carry out normal cellular functions. In addition to well-established actin cytoskeleton regulators, such as nucleators, capping proteins, and bundlers, cells likely have uncharacterized modulators that regulate cytoskeleton dynamics, the detailed functions of which are not yet fully understood. In this study, we conducted biochemical exploration to identify the actin-regulatory activity of Kaptin (KPTN), a protein known to co-localize with actin-rich structures at the cell’s periphery. Using single-molecule assays, we demonstrated that KPTN inhibits actin nucleation. Our results revealed that KPTN possesses a novel barbed-end capping activity, which stabilizes and bundles actin filaments. Structural modeling, based on AlphaFold, suggests that KPTN is a member of the WD-repeat-containing protein family. Furthermore, we identified a crucial cationic residue in the putative N-terminal beta-propeller region of KPTN that plays a critical role in modulating actin dynamics. In summary, our data unveil the mechanistic underpinning functions of KPTN and establish its novel role as a regulator of the actin cytoskeleton.

## Introduction

Cellular processes like cell motility, intracellular transport, and endocytosis rely on the dynamic remodeling of the actin filament-based structures including lamellipodia, filopodia, microvilli, and stress fibres (Pollard and Cooper 1986; Insall and Machesky 2009; Pollard and Cooper 2009). Actin filament dynamics are regulated by multiple actin-associated proteins at various levels, including nucleation, elongation, capping, crosslinking, disassembly, and turnover of the actin filaments (Campellone and Welch 2010; Dominguez 2009; Chesarone et al., 2010).

The reorganization of actin at the leading edge is tightly governed in a spatiotemporal manner by actin-binding proteins to produce filaments of specific lengths (Bugyi and Carlier 2010). Short-branched filaments are predominant in lamellipodia whereas long unbranched filaments are prevalent in filopodia (Fujiwara et al., 2014; Breitsprecher and Goode 2013). Short-branched filaments in lamellipodia are predominantly modulated by the plus end capping protein (CapZ) and Arp2/3 complex, while actin nucleators like formins and elongator Ena/VASP modulate the long unbranched filaments in filopodia (Funk et al., 2021; Wear et al., 2003; Chesarone et al., 2010; Goode and Eck, 2007; Kovar et al., 2006; Rottner et al., 2017; Zweifel et al., 2021).

Kaptin (KPTN) was identified as an actin-associated protein in a screen for actin-binding proteins in platelets using F-actin affinity chromatography (Bearer 1992). Initial studies revealed KPTN as a novel actin-binding protein localized at the leading edge of the migrating cells like lamellipodia of chick fibroblast cells, growth cone filopodial of PC-12 cells, and at the tip of the stereocilia of the hair cells of the inner ear (Bearer 1992a; Bearer et al., 2002; Bearer and Abraham 1999; Bearer et al., 2000). KPTN was mapped to chromosome 19q13.4 by performing sequencing of the cloned cDNA from HEL cells and it was suggested to be a candidate gene for the DFNA4 hearing loss locus (Bearer et al., 2000). Recently, a pathogenic homozygous variant of the KPTN gene expressed in neuronal cells and stereocilia cells was identified in individuals with hearing impairment (Liaqat et al., 2023). Initial analysis of the KPTN protein sequence did not reveal any known protein domain that could provide insights into its functions within the cells (Bearer and Abraham 1999; Baple et al., 2014).

In recent times, KPTN has been discovered as a component of the KICSTOR complex which includes KPTN, ITFG2, C12orf66, and SZT2, and is involved in regulating the mTORC1 signaling (Wolfson et al., 2017). These megadalton KICSTOR complex components are conserved in vertebrates but not in non-vertebrates and fungi (Wolfson et al., 2017). In cultured cells, the gene products of the KICSTOR complex are necessary to inhibit the mTORC1 pathway in response to amino acid or glucose deprivation (Wolfson et al., 2017).

KPTN-knockout mice and cortical neural precursors differentiated from human iPSCs lacking KPTN display transcriptional and biochemical evidence of altered mTOR signaling, supporting KPTN’s role in mTORC1 regulation (Levitin et al., 2023). Mutations in SZT2 or C12orf66 have been associated with brain malformation disorder (Basel-Vanagaite et al., 2013; Venkatesan et al., 2016; Mc Cormack et al., 2015). Mutations in the KPTN gene have been linked to human macrocephaly, neurodevelopmental delay, and moderate to severe epilepsy (Baple et al., 2014; Pajusalu S et al., 2015; Thiffault I et al., 2018; Miguez MP et al., 2020; Horn S et al., 2023, Ullah A et al., 2022).

An immunofluorescence study on rat hippocampal neurons revealed the localization of endogenous and GFP-tagged wild-type KPTN to actin-rich structures in neurons (Baple et al., 2014). This localization was lost in the KPTN protein carrying pathogenic mutations (Baple et al., 2014). Despite multiple lines of evidence suggesting a role for KPTN in actin remodelling, a detailed mechanistic understanding of KPTN as an actin modulator is lacking.

In this study, we identify KPTN as a member of the WD-repeat-containing protein family with a predicted 7-bladed beta-propeller structure. Using conventional biochemical approaches and TIRF microscopy, we demonstrate that KPTN inhibits the elongation of barbed end actin filaments. Furthermore, we show that KPTN stabilizes actin filaments and forms thin actin bundles. We also find that a key residue within the predicted β-propeller region of KPTN mediates interactions with actin and is necessary for inhibiting actin filament elongation. Our work represents the first report to describe the biochemical function of KPTN as a weak barbed end-capping protein, emphasizing its role as an actin cytoskeleton modulator.

## Result

### 2.1 KPTN binds to actin filaments

In previous studies, KPTN was purified from platelet cell lysates using an F-actin affinity column (Bearer 1992a). The presence of KPTN at the edge of the lamellipodia of fibroblast cells suggested its role in actin dynamics (Bearer 2000). To assess the role of KPTN in lamellipodial actin dynamics, we investigated the biochemical function of KPTN in regulating actin filament turnover.

We analyzed the full-length human KPTN (hKPTN) (1-436) amino acid sequence (Fig 1A) and found a predicted disordered region in the C-terminal end (amino acids 421-436) (Fig 1B). The (1-420) amino acids of hKPTN were cloned into pSNAP vector (NEB) and purified using the 6X-His-based affinity system (Fig 1C, Fig 1D).

**Figure 1:**
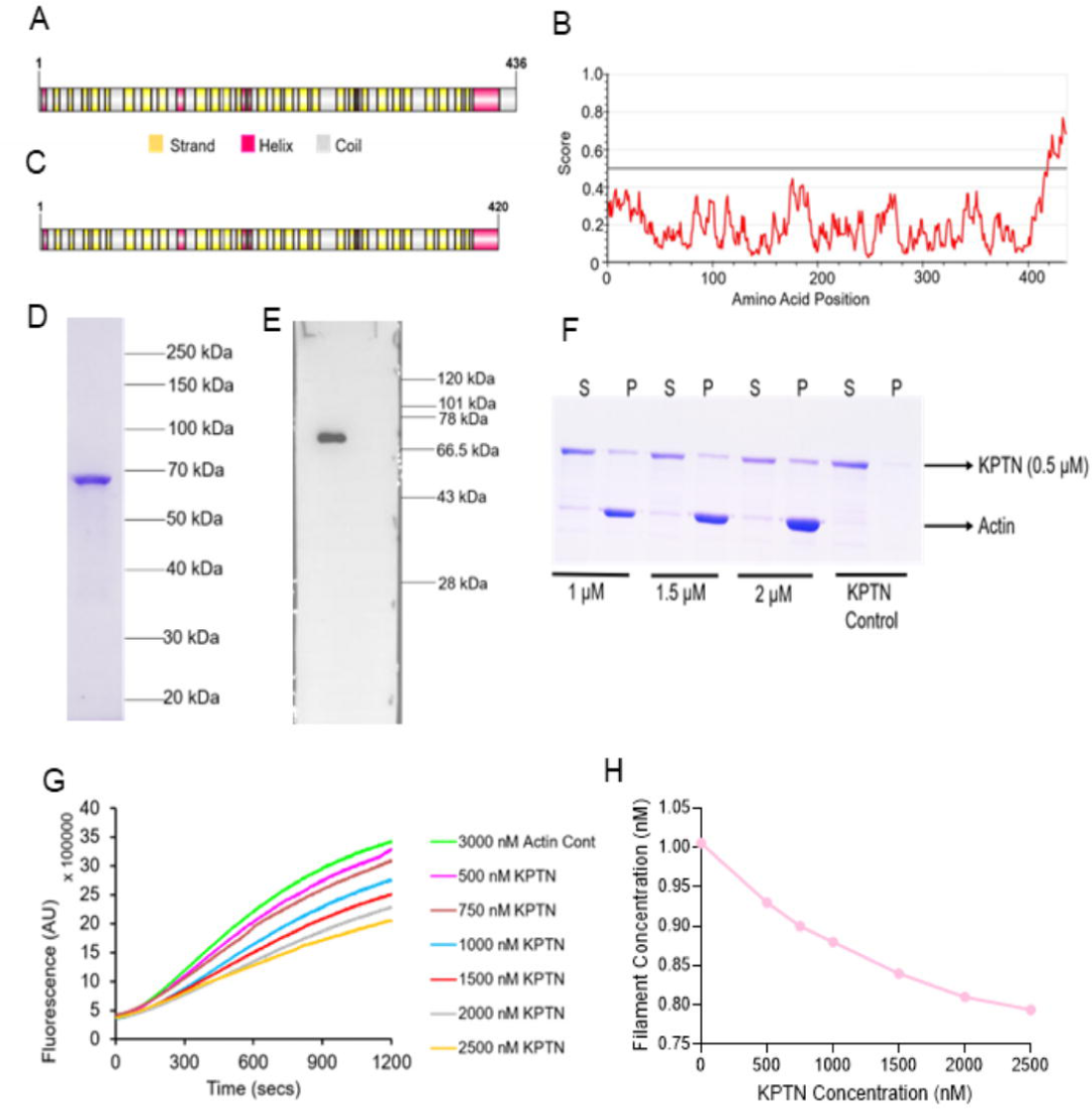
Human KPTN binds actin filaments. (A) Schematic representation of full-length human Kaptin (hKPTN) with predicted secondary structure. (B) IUPred2 identifies the disordered region in KPTN (421-436 aa) [> 0.5 considered as a disordered region in the protein]. (C) The [1-420] amino acid region was used for this study. (D) 10% SDS PAGE showing the SNAP-tag hKPTN 6X-His tag protein purified using a bacterial expression system. (E) Western Blot was performed using the commercially available Sigma antibody to detect the purified hKPTN protein. (F) High-speed (320,000 g) co-sedimentation assay of 1-420 region of hKPTN to actin filaments. An increasing concentration of actin filaments was incubated with 2 µM hKPTN. S and P represent the supernatant and pellet respectively. The data was repeated independently five times. (G) Time-course of pyrene-actin nucleation assembly assay with the indicated concentrations of the hKPTN. 3000 nM actin (10% pyrene-labelled) and indicated (nM) concentrations of hKPTN protein were used. The data was repeated independently five times. (H) The plot of filament concentration in between 300-400 seconds as a function of various concentrations of hKPTN.

The purified hKPTN was subjected to the Superose 6 (10/300 GL) column for further protein purification (Fig S1A). We obtained a single-band purified hKPTN protein through this FPLC-based purification (Fig 1D). Furthermore, we confirmed the purified hKPTN by western blotting using the anti-KPTN antibody (Sigma) (Fig 1E). Mass spectrometry was performed on the in-gel trypsin-digested purified protein validating the presence of hKPTN (Fig S1B and S1C). To directly probe the interaction of hKPTN with actin filaments, we performed the conventional high-speed co-sedimentation assay with purified hKPTN protein (Dutta et al., 2017; Kundu et al., 2021). The co-sedimentation data revealed that hKPTN co-sediments with actin filaments in a concentration-dependent manner (Fig 1F). hKPTN remains in the supernatant fraction when no actin filaments were provided (Fig 1F). These data demonstrate the successful *in vitro* purification of hKPTN and its binding to actin filaments.

### 2.2 hKPTN inhibits actin assembly

As hKPTN exhibited binding with actin filaments, we analyzed the activity of the purified hKPTN protein using *in vitro* pyrene-actin assembly (Dutta et al., 2017, Dutta et al., 2019). In fluorometric nucleation assays, hKPTN was observed to inhibit the actin assembly in a dose-dependent manner (Fig 1G). We calculated the filament concentration using the slopes between 300-400 seconds of polymerization assuming an elongation rate of 3.62 µm ^-1^ sec ^-1^ based on rates of 11.16 and 1.3 µm ^-1^ sec ^-1^ for the barbed end and pointed end respectively (Fig 1H) (Harris ES et al., 2004). We observed that with increasing concentration of hKPTN, the amount of filament formed was reduced (Fig 1H). Previous studies with chick embryo fibroblast and PC-12 cell lysates, and purified actin filaments indicated reduced actin filament binding to the KPTN-antibody coated coverslips in the presence of KPTN (Bearer 1992a). Control data with only antibody-coated coverslips showed no binding of actin filaments to the coverslips. This data is consistent with our results showing KPTN prevents actin assembly.

Total internal reflection fluorescence (TIRF) microscopy was conducted with different concentrations on hKPTN and 1000 nM of actin monomers. In the presence of hKPTN, actin filaments were shorter and sparser compared to actin-only controls (Fig 2A-D, and S2A) and exhibited a concentration-dependent reduction in filament number and growth (Fig 2E-F). The calculated mean length of actin filaments in actin-only controls at 0 seconds was 6.21 ± 0.3 (SEM) µm while the mean actin filament length in the presence of 2500 nM hKPTN is 3.13 ± 0.22 (SEM) µm (Fig S2B). When the filament length was measured at 480 seconds, the mean actin filament length was 11.72 ± 0.42 (SEM) µm for actin-only controls and 5.72 ± 0.23 (SEM) µm for 2500 nM hKPTN (Fig S2B). The elongation rate of the filament was calculated, considering 1 actin subunit contributes 2.7 nm to the actin filament length (Hoeprich et al., 2022, Breitsprecher et al., 2012; Henty-Ridilla 2022). The TIRF microscope imaging began with actin filaments having a few microns of length at the first frame, and the subsequent elongation was evaluated (Fig S2C). The mean elongation rate for actin-only controls was 13.03 ± 0.33 (SEM) subunits µM ^-1^ sec ^-1^ whereas it was 5.63 ± 0.19 (SEM) subunits µM ^-1^ sec ^-1^ for 2500 nM hKPTN-mediated reaction (P < 0.0001) (Fig 2F). To understand the affinity of hKPTN for the barbed end, we calculated the K_d_ of hKPTN by plotting the fraction of free-growing ends versus the hKPTN concentration (Fig 2G). The binding affinity of hKPTN to the actin filaments is 758.6 nM.

**Figure 2:**
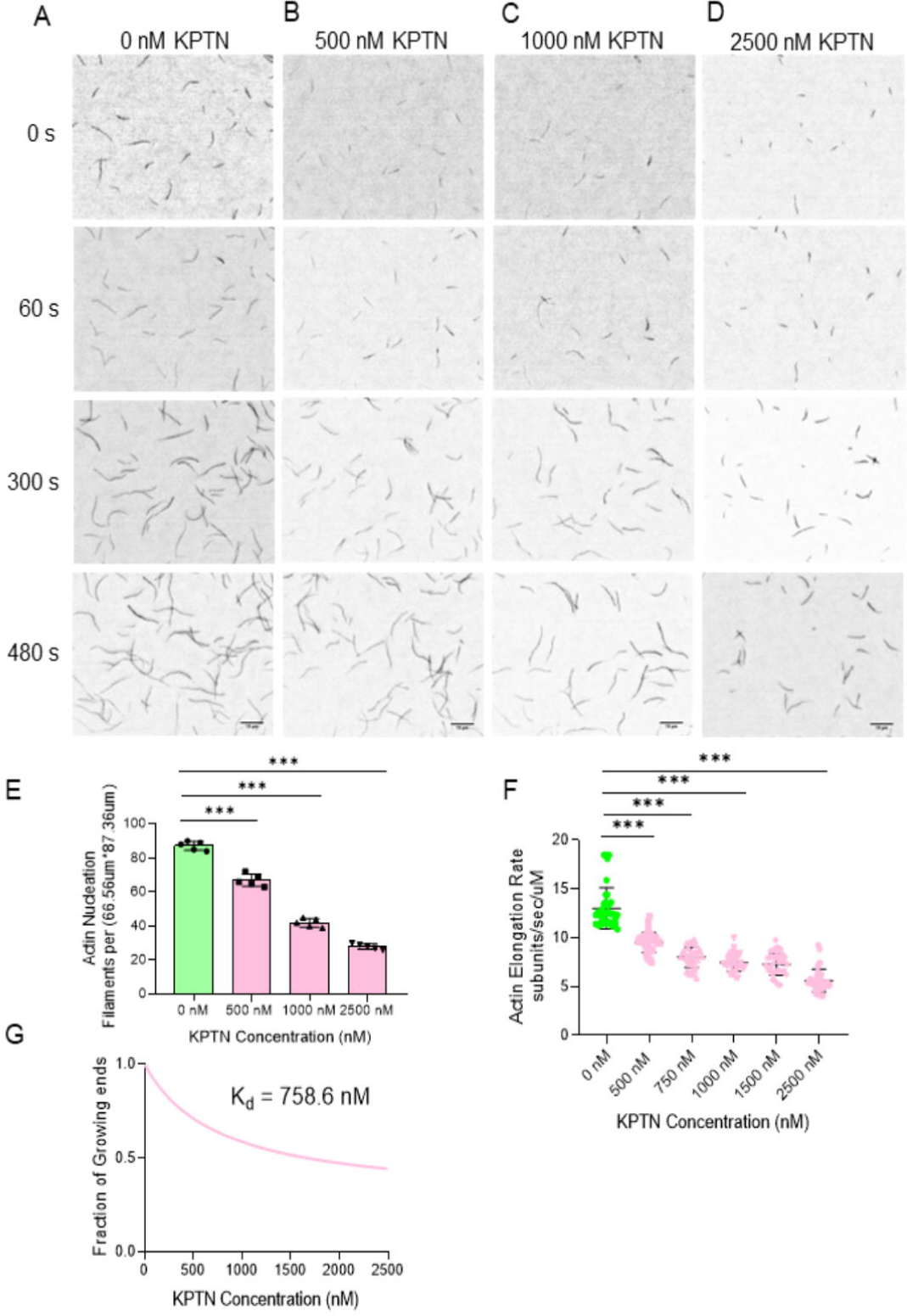
hKPTN inhibits actin assembly. Time-lapse TIRF microscopy images of the formation of actin filaments from 1000 nM actin monomers in the presence of (A) 0 nM hKPTN, (B) 500 nM hKPTN, (C) 1000 nM hKPTN and (D) 2500 nM hKPTN. Scale bar 10 µm. Imaging was started after 90 seconds of keeping the reaction inside the flow chamber. (E) The average number of filaments was measured at 480 seconds for 0 nM, 500 nM, 1000 nM, and 2500 nM of hKPTN concentrations. The mean and SEM represent the data from three independent experiments (n=3 field of view) ***P<0.0001 One-way non-parametric ANOVA was performed. (F) The actin filament elongation rate (subunit/sec/µM) was calculated assuming 370 subunits per micron of actin filaments. The filaments observed from the first frame were considered for analysis. Minimum (n=40) actin filaments in each case were taken for the analysis. ***P<0.0001 Unpaired t-tests were done to analyze the data. (G) The fraction of free-growing ends versus the concentration of hKPTN was plotted using the hyperbolic curve to determine the binding affinity of hKPTN which is 758.6 nM for the actin filaments.

With a high concentration (3000 nM) of actin monomers, hKPTN constrained actin filament assembly (Fig S3A-C). The inhibitory effect with a high concentration of hKPTN was pronounced, but the filaments could not be quantified. We also observed thin bundles of actin filaments, but the actin filament length could not be measured (Fig S3D). Together these results indicate that hKPTN binds to actin and prevents actin assembly.

### 2.3 hKPTN bundles actin filament

During actin filament assembly with a high concentration of actin monomers, we observed the formation of thin actin bundles in the presence of hKPTN (Fig S3). To elucidate the bundling activity of hKPTN, we evaluated actin filaments at low concentrations in the absence and presence of hKPTN using the TIRF microscope. Actin filaments by themselves did not form bundles (Fig S4A and S4C). Strikingly, in the presence of hKPTN, we observed the formation of thin actin bundles (Fig S4B and S4C). These reactions were further examined using a high-resolution transmission electron microscope (TEM). The TEM images show no actin bundles in the absence of hKPTN, but actin filament bundles were closely observed in the presence of hKPTN (Fig S4D). Based on these data, we suggest that hKPTN acts as a weak bundling protein that inhibits actin assembly.

### 2.4 hKPTN does not promote barbed-end actin elongation

KPTN-mediated hindrance of actin filament growth prompted us to analyze KPTN’s ability to elongate actin filaments using the seeded actin assembly assay. Earlier studies speculated that KPTN was a barbed end-binding protein (Bearer EL 1992a; Bearer EL et al., 2002). We prepared 333 nM of filamentous actin seeds by shearing pre-formed actin filaments (Moseley et al., 2006). In the presence of 400 nM actin monomers (to inhibit monomer incorporation into the pointed end of the filament), actin filaments in the actin-only control reaction were observed to grow (Fig 3A) while no growth of the actin filaments was observed when the barbed-end capping protein CapZ was included in the reaction mixture (Fig 3B). The addition of hKPTN in seeded actin assembly resulted in negligible to no growth of the actin filaments in a dose-dependent manner as compared to the actin control without hKPTN and CapZ (Fig 3C and 3D). Therefore, this data suggests hKPTN inhibits barbed-end actin filament elongation. The mean filament length was calculated at 0 seconds and after 480 seconds. At 0 seconds, the mean filament length for 0 nM hKPTN (i.e., actin-only control) was 3.139 ± 0.17 µm (SEM) and in the presence of 600 nM, hKPTN was 2.47 ± 0.08 µm (SEM). At 480 seconds, the mean filament length for actin-only control was 7.27 ± 0.37 µm (SEM) and for 600 nM hKPTN was 2.99 ± 0.12 µm (SEM) (P < 0.0001) (Fig 3E). Our data suggests that a high concentration of hKPTN does not accelerate the barbed-end elongation of actin filaments.

**Figure 3:**
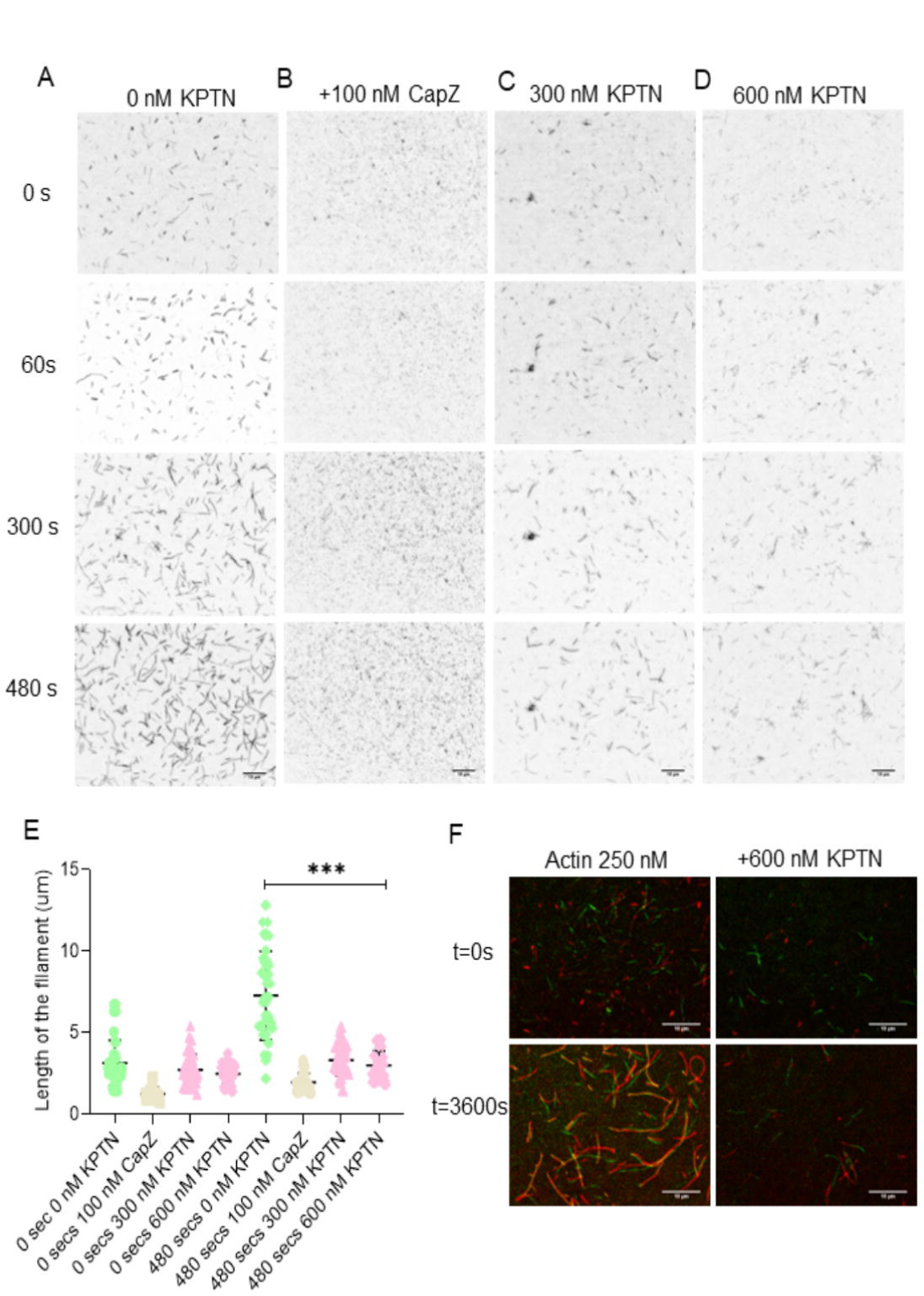
hKPTN caps the barbed end of the filament. Time-lapse TIRF microscopy images of seeded actin assembly with 333 nM actin seeds and 400 nM actin monomers (barbed end kinetics). Time-lapse images from TIRF microscopy using seeded actin assembly with (A) 0 nM hKPTN (B) 100 nM CapZ, (C) 300 nM hKPTN, and (D) 600 nM hKPTN. Imaging was started after 40 seconds of the reaction mixture in the flow chamber. Scale Bar 10 µm. (E) The average length of the filaments was measured at the beginning (0 seconds) and end (480 seconds) of imaging for 0 nM KPTN (n= 55), 100 nM CapZ (n=30), 300 nM KPTN (n= 40), and 600 nM hKPTN (n=50). The experiment was repeated thrice. The error bar represents the SEM. ***P<0.0001 One-way non-parametric ANOVA was performed. (G) Direct visualization of filament reannealing in the absence and presence of barbed end capping protein. Equimolar concentration i.e., 250 nM of dual colour (rhodamine phalloidin and Alexa-488 phalloidin stabilized) labeled actin filaments were sheared with a 27-gauge needle and imaged immediately within 40 seconds and after 3600 seconds. hKPTN inhibits the reannealing of the actin filaments. Scale Bar 10 µm.

To evaluate the effect of hKPTN on the barbed-end kinetics of the actin filaments, we performed the actin filament reannealing assay using dual-colour phalloidin stabilized actin filaments (Andrianantoandro et al., 2001; Patel et al., 2018). 250 nM of actin filaments were sheared with a 27-gauge needle and allowed to reanneal (Andrianantoandro et al., 2001; Patel et al., 2018). After 60 minutes, we observed that in the absence of hKPTN, actin filaments could reanneal. Unlike the actin-only control, hKPTN inhibited actin filament reannealing, indicating that it binds to the barbed end and prevents filament annealing (Fig 3F). This observation is consistent with our previous result, demonstrating hKPTN binds to the barbed end of the actin filament and restricts the actin filament elongation.

### 2.5 Effect of Profilin in seeded actin elongation in the presence of hKPTN

Profilin binds to the barbed side of actin, which is available on both G-actin and F-actin at the filament barbed end. Profilin enhances the barbed-end kinetics and modulates the actin filament length (Pernier et al., 2016). Profilin plays an important role in providing ATP-bound actin monomers to the growing ends of the actin filament (Selden et al., 1999). Therefore, we performed the seeded filament assembly in the presence of profilin under the TIRF microscope (Fig 4A). The inhibitory effect of hKPTN was reduced in the presence of profilin compared to the reaction without profilin (Fig 4B, 3C-D). The mean length of the actin filament at 480 secs in the presence of profilin but without hKPTN is 7.46 ± 0.24 µm (SEM) is similar to the actin filament growth in the absence of both hKPTN and profilin (Fig 4C and 3E). When hKPTN is combined with profilin, it allows the actin filament growth, and the mean length of the actin filament at 480 seconds is 5.81 ± 0.29 µm (SEM) (Fig 4C). These results demonstrate that hKPTN permits the barbed end actin dynamics in the presence of profilin.

**Figure 4:**
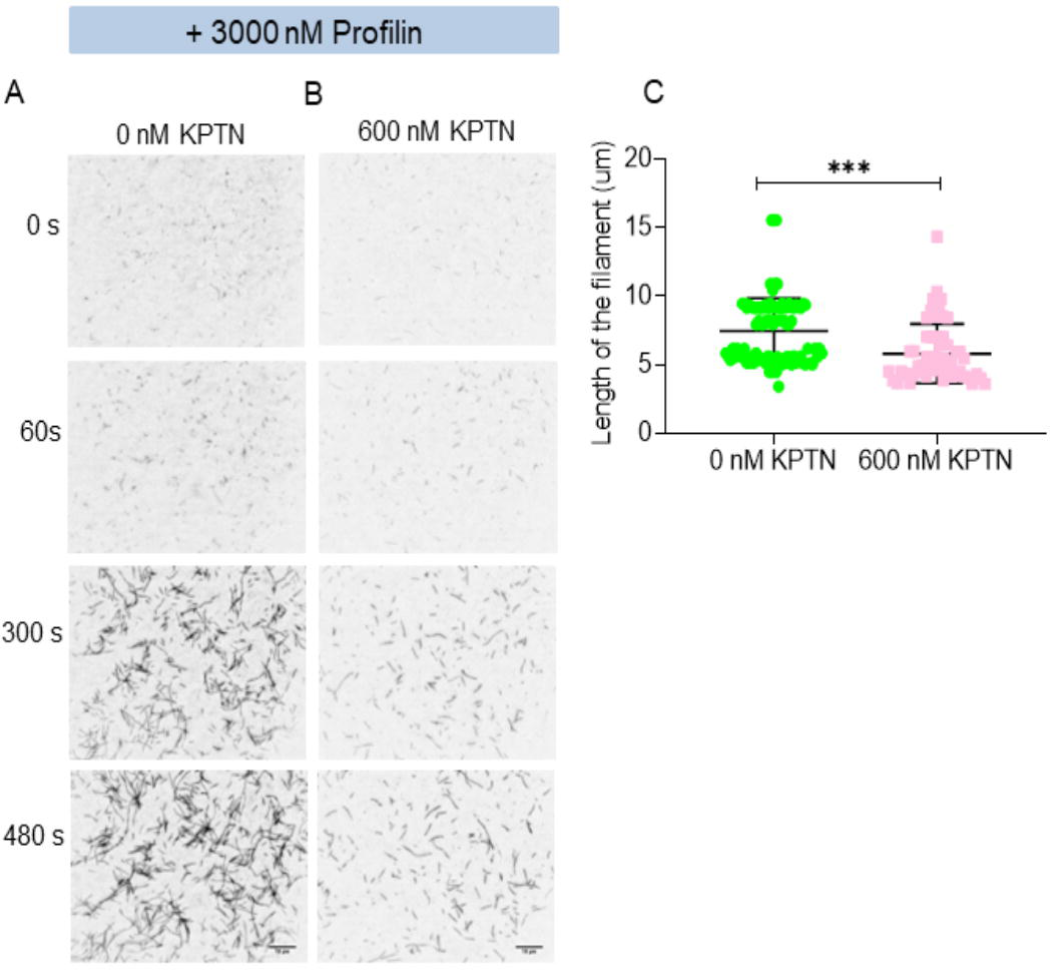
Profilin inhibits the capping activity of hKPTN. (A) Time-lapse TIRF microscopy images of seeded actin assembly with 333 nM actin seeds and 400 nM actin monomers (barbed end kinetics) in the presence of 3000 nM Profilin and 0 nM hKPTN. (B) Time-Lapse TIRF microscopy images of seeded actin assembly with 400 nM actin monomers, 3000 nM Profilin, and 600 nM hKPTN. Imaging was started after 40 seconds of the reaction mixture in the flow chamber. Scale Bar 10 µm. (C) The length of the actin filaments was measured in the presence of Profilin without hKPTN (n= 95) and with KPTN (n= 54). The experiment was repeated thrice. The error bar represents the SEM. ***P<0.0001 unpaired T-Test was performed.

Previous studies have shown that KPTN elutes from the F-actin affinity column in the presence of ATP (Bearer 1992a; Bearer et al., 2002). Consistent with this observation, our results suggest that hKPTN facilitates actin elongation when profilin promotes ATP-bound actin monomer addition to the barbed end.

### 2.6 hKPTN inhibits actin filament depolymerization

Our current studies suggest that hKPTN acts as a barbed end-capping protein. However, the capping activity of hKPTN is not as robust as that of the canonical capping protein CapZ (Fig 3). hKPTN allows the slow growth of actin filament in the presence of profilin (Fig 4). Further, we investigated the depolymerization kinetics using positive (Cofilin) and negative (CapZ) controls to understand the effect of hKPTN on actin filament depolymerization. We diluted 5000 nM actin filaments to 100 nM to carry out the dilution-dependent actin depolymerization kinetic studies. Actin filament depolymerization was inhibited and filaments were stabilized in the presence of hKPTN (Fig 5E and 5F) similar to filament stabilization in the presence of barbed end capping protein CapZ (Fig 5C and 5D). In the case of actin-only control subjected to actin depolymerization assay, the majority of actin filaments had a length of up to 4 µm (Fig 5G). Interestingly, for hKPTN most of the actin filaments had a length not exceeding 12 µm whereas the majority of the CapZ-treated actin filaments had a length of 16 µm (Fig 5G). From these data, we infer that similar to CapZ, hKPTN also binds to the barbed end of the actin filaments thereby inhibiting actin filaments depolymerization and stabilizing them.

**Figure 5:**
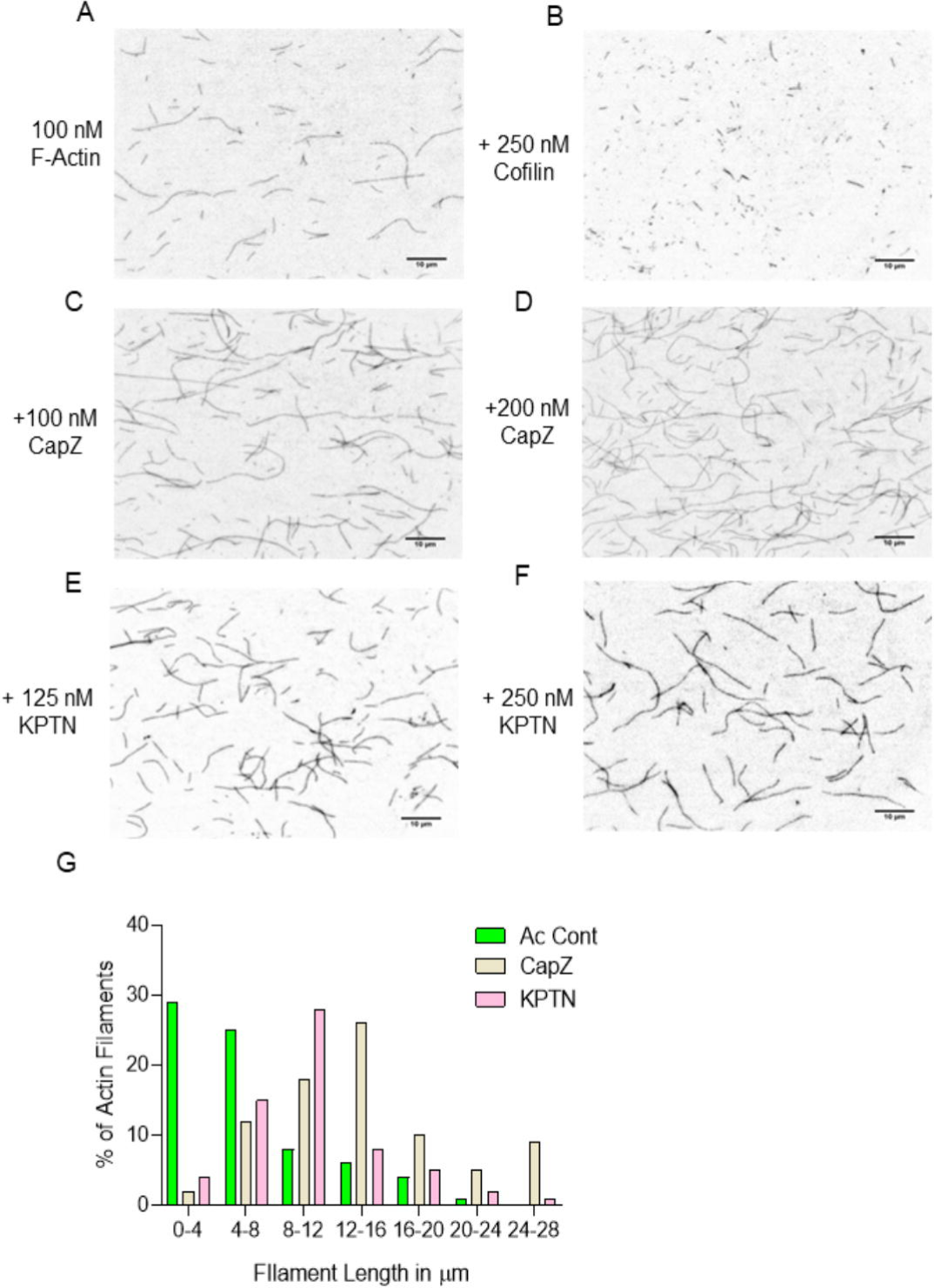
hKPTN stabilizes actin filaments. (A) 5000 nM of actin filaments was diluted to a final concentration of 100 nM. Visualization of 100 nM actin filaments depolymerization. The images were recorded after 120 seconds of the reaction mixture in the flow chamber. Dilution-dependent depolymerization of actin filaments in the presence of (B) 250 nM Cofilin, (C) 100 nM CapZ, (D) 200 nM CapZ, (E) 125 nM hKPTN and (F) 250 nM hKPTN. (G) Filament length distribution for filaments in buffer (green colour-actin), 100 nM CapZ (yellow colour), and 125 nM hKPTN (pink colour). Most of the filaments for actin control have lengths between 0-4 µm. In the case of 100 nM CapZ, maximum filaments have lengths in between 12-16 µm. For 125 nM hKPTN, most filaments have lengths below 12 µm. The experiment was repeated for five times.

### 2.7 hKPTN binds to the barbed end of the actin filament

We were curious to visualize the localization of hKPTN on the actin filament and the growth of actin filaments in the presence of hKPTN. To address this, we generated fluorescently labeled SNAP-tag hKPTN construct following the standard protocol (Breitsprecher et al., 2012). The fluorescent labeling percentage of the hKPTN was between 30%-40% as calculated following standard procedure (Breitsprecher et al., 2012). In the reaction mixture with preformed actin filaments, we could see the localization of the fluorescently labeled SNAP-tag hKPTN on the actin filaments and some free SNAP-labeled hKPTN in the field (Fig 6A). The fluorescently labeled hKPTN was localized to the tip of the actin filament.

**Figure 6:**
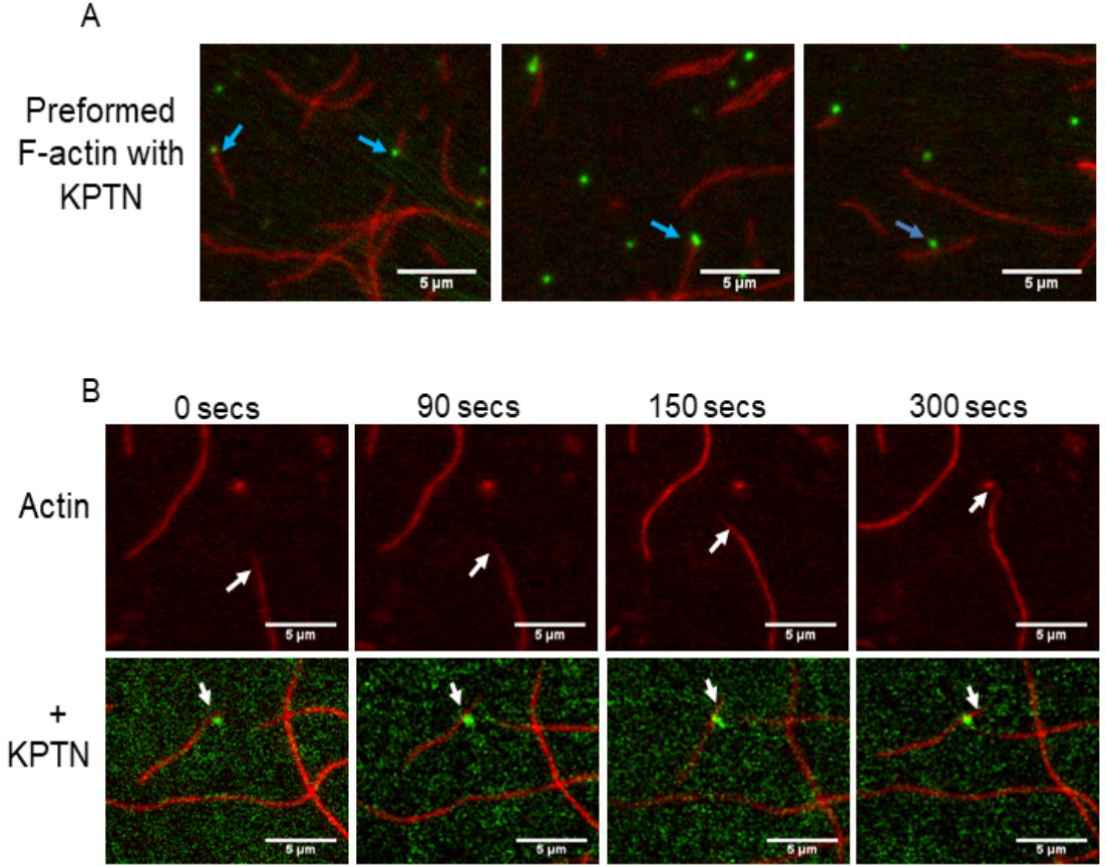
hKPTN present at the tip of the filament. (A) The different view of the KPTN localization on the actin filament. The preformed actin filament along with 30%-40% labelled SNAP-tag KPTN showed the presence of hKPTN at the tip of the filament. (B) Time-lapse TIRF microscopy images of actin filaments with fluorescently labeled hKPTN. The actin filament without KPTN is growing freely but hKPTN-mediated actin filament growth is inhibited. Imaging was started after 40 seconds of the reaction mixture in the flow chamber. Scale Bar 5 µm. The experiment was repeated thrice.

To understand the effect of hKPTN on actin filament growth, we conducted actin filament assembly experiment both in the presence and absence of fluorescently labeled hKPTN. Our observations revealed that the growth of actin filaments was impeded in the presence of hKPTN, compared to when hKPTN was absent from the reaction mixture (Fig 6B). This data, in line with the results presented in the previous section, led us to infer that hKPTN acts as a capping protein that decelerates the growth of actin filaments.

### 2.8 KPTN is a member of the WD-repeat-containing protein family

Recently the AlphaFold protein structure database predicts the hKPTN protein structure comprised of an N-terminal 7-bladed β-propeller domain followed by a C-terminal coiled-coil domain and an unstructured loop structure (Fig S5A and Fig S5A’) (Jumper et al., 2021; Tunyasuvunakool et al., 2021). The boundaries of each blade in the β-propeller as speculated by AlphaFold are displayed in Fig S5B. As hKPTN was predicted to have a 7-bladed β-propeller structure, we performed the sequence-based phylogenetic analysis of hKPTN and the well-characterized actin-binding proteins with a 7-bladed β-propeller structure. For the phylogenetic analysis, we considered the sequence of vertebrates because KPTN being a member of the KICSTOR family is well characterized in higher eukaryotes. The phylogenetic analysis revealed KPTN to be a nearby member of the Coronin family WD-repeat-containing proteins (Fig S5C). The protein sequence of Coronin and KPTN were aligned using Clustal Omega (Fig S6). Coronin belongs to the well-characterized WD-repeat-containing protein family (Appleton BA et al., 2006). From the protein sequences, we recognized three WD-repeats in Coronin-1A, two WD-repeats in Coronin-1B, and one WD-repeat in Coronin-1C. The sequence analysis revealed that the mouse KPTN has one WD-repeat and in hKPTN the aspartic acid remained conserved but the tryptophan was modified to arginine (Fig S6). Similarly, in the case of Coronin-1C, the tryptophan amino acid is conserved, and the aspartic acid was modified to asparagine residue (Fig S6). For the first time, this result implicates KPTN protein as a member of the WD-repeat-containing protein family along with other structurally conserved actin-binding proteins.

### 2.9 A conserved Arginine residue in the beta-propeller region of KPTN is involved in actin-binding

Studies with mammalian Coronin have identified a conserved amino acid residue (Arg^30^) within the N-terminal β-propeller region which when mutated to aspartic acid (R30D) reduced the actin filament binding activity (Cai et al., 2007). Alignment of vertebrate Coronin and KPTN protein sequence identified hKPTN Arg^59^ to be equivalent to Arg^30^ of Coronin (Fig. S6). This Arg^59^ residue in hKPTN is also present in the β-propeller region (Fig S5 A’’). The positively charged residue arginine (Arg^59^ in hKPTN) is conserved for the higher eukaryotes except for *Danio rerio* (Fig S7A) where the position of positively charged (arginine) has been interchanged with the neighbouring neutral glutamine (Fig. S7B). Our study highlights the residue in KPTN which is conserved and probably modulates actin dynamics. This work is the initial beginning of underpinning the actin-binding sites in the KPTN protein.

We mutated (Arg^59^) to aspartic acid residue in the wild-type hKPTN protein based on our analysis above and according to the available information on the mammalian Coronin actin-binding site (Cai et al., 2007; Gandhi et al.,2010; Chan et al., 2012). The hKPTN^(R59D)^ (1-420) was purified like the wild-type hKPTN [hKPTN^(WT)^] (Fig 7A). Actin co-sedimentation assay was used to investigate the actin-binding of hKPTN^(R59D)^. Our data revealed that the R59D mutation abolished the actin-binding activity of hKPTN (Fig 7B). This data is consistent with the R29D and R30D mutants of Coronin-1A and 1B respectively (Chan et al., 2012).

**Figure 7:**
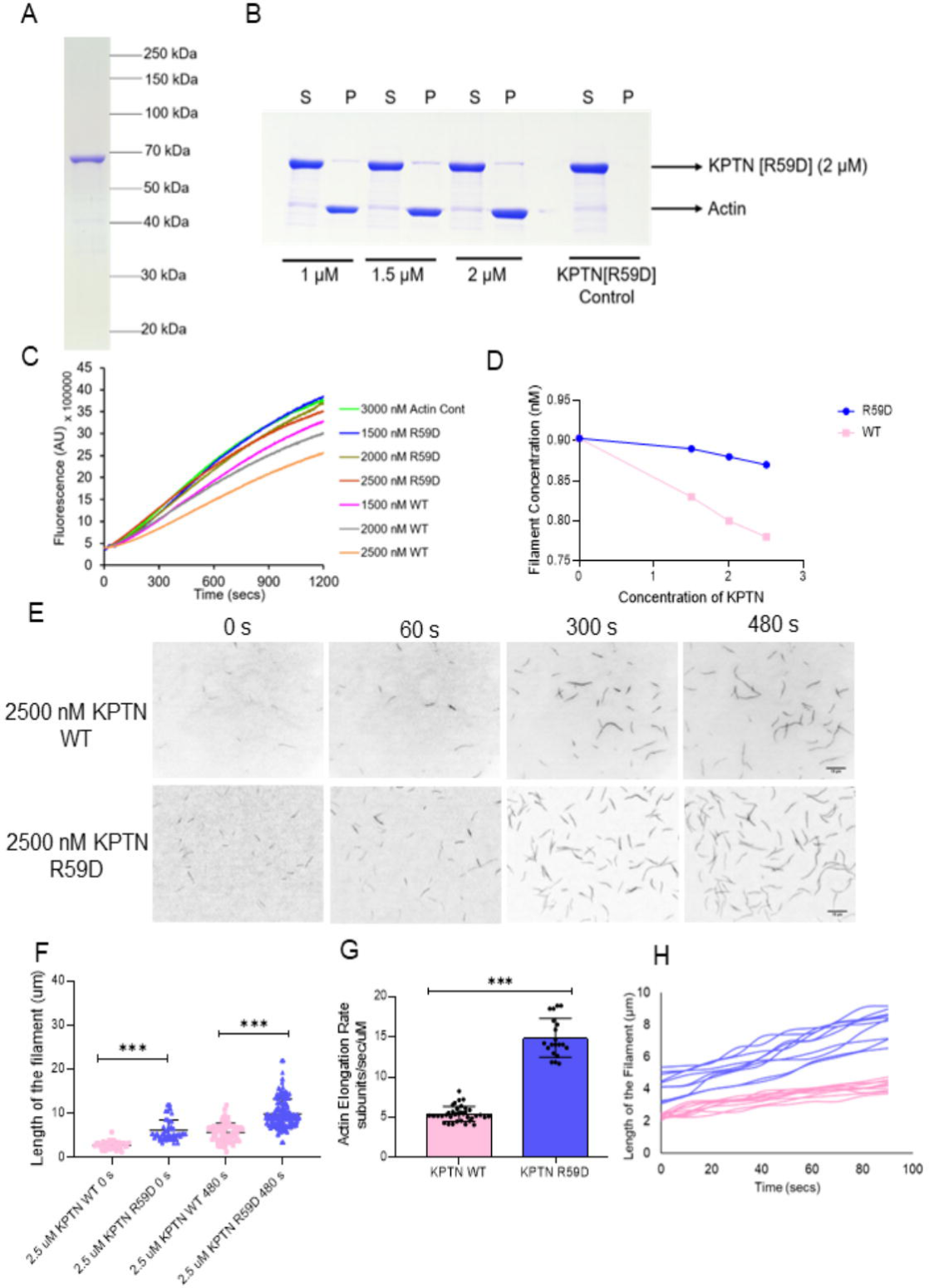
hKPTN^(R59D)^ mutant abolishes inhibition of actin assembly. (A) 10% SDS PAGE showing the SNAP-tag R59D hKPTN protein purified using a bacterial expression system. (B) High-speed (320,000 g) co-sedimentation assay of 1-420 region of R59D hKPTN to actin filaments. An increasing concentration of actin filaments was incubated with 2 µM R59D hKPTN. S and P represent the supernatant and pellet respectively. The experiment was performed thrice. (C) Time-course of pyrene-actin nucleation assembly assay with the indicated concentrations of the hKPTN^(R59D)^ and hKPTN^(WT)^. 3 μM actin (10% pyrene-labeled) and indicated (nM) concentrations of the protein were used. The experiment was performed five time independently. (D) The plot of filament concentration in between 300-400 seconds as a function of various concentrations of hKPTN^(R59D)^ and hKPTN^(WT)^. (E) Time-lapse imaging of 2500 nM of hKPTN^(R59D)^ and hKPTN^(WT)^ along with 1000 nM actin monomers. (F) The average length of the filaments was measured at different time points for 2500 nM hKPTN^(R59D)^ and hKPTN^(WT)^. The mean and SEM represents the data from three independent experiments. ***P<0.0001 One-way non-parametric ANOVA was performed. (G) Actin filament elongation rate (subunit/sec/µM) was calculated assuming 370 subunits per micron of actin filaments. The filaments observed from the first frame were considered for analysis. (n=37) hKPTN^(WT)^ and (n=25) hKPTN^(R59D)^ actin filaments were taken for analysis. ***P<0.0001 Unpaired t-tests were done to analyze the data. (H) Length of the actin filaments over time plotted for hKPTN^(WT)^ [pink] and hKPTN^(R59D)^ [blue].

Our study is the first ever to characterize the actin-binding site on the hKPTN protein. Further, we tested the hKPTN^(R59D)^ for bulk polymerization assay. The hKPTN^(R59D)^ mutant in comparison to wild-type hKPTN^(WT)^ showed negligible to no inhibition of actin polymerization in the *in vitro* pyrene actin assembly (Fig 7C). We quantified the filament concentration for the hKPTN^(R59D)^ following the methods described in the earlier section. Our result does not reflect any significant change in the filament concentration for the hKPTN^(R59D)^ from the actin-only control while hKPTN^(WT)^ showed decreased filament concentration with increasing concentration of hKPTN (Fig 7D).

TIRF microscopy was performed with 1000 nM actin monomers for hKPTN^(R59D)^ and revealed no inhibition in actin assembly in comparison to the hKPTN^(WT)^ (Fig 7E). The average length of the actin filaments for KPTN^(WT)^ at 0 seconds is 2.78 ± 0.14 (SEM) µm and for hKPTN^(R59D)^ is 6.1 ± 0.29 (SEM) µm (Fig 7F). Similarly, at 480 seconds, the average length of the actin filaments for hKPTN^(WT)^ is 5.67 ± 0.24 (SEM) µm, and for hKPTN^(R59D)^ is 9.89 ± 0.23 (SEM) µm (Fig 7F). The actin filament elongation rate for the hKPTN^(R59D)^ is 14.88 ± 0.54 (SEM) subunits µM ^-1^ sec ^-1^ in contrast to the hKPTN^(WT)^ has an elongation rate of 5.43 ± 0.15 (SEM) subunits µM ^-1^ sec ^-1^ (Fig 7G and 7H). These observations indicate that the (Arg^59^) position of hKPTN binds to actin filament and plays an important role in actin cytoskeleton regulation by hKPTN.

### 2.10 R59D hKPTN does not inhibit seeded actin filament elongation

The seeded actin assembly was carried out as indicated in the earlier section of this study with 333 nM actin seeds and 400 nM actin monomers. The actin filaments elongate without any inhibition in the reaction mixture without the hKPTN^(WT)^construct (Fig 8A). Our hKPTN^(R59D)^ construct does not restrain the actin elongation (Fig 8B). The length of the actin filaments after 480 seconds in the case of the hKPTN^(R59D)^ construct is 8.145 ± 0.12 (SEM) µm, similar to the reaction in the absence of hKPTN^(R59D)^ which is 8.54 ± 0.36 (SEM) µm.

**Figure 8:**
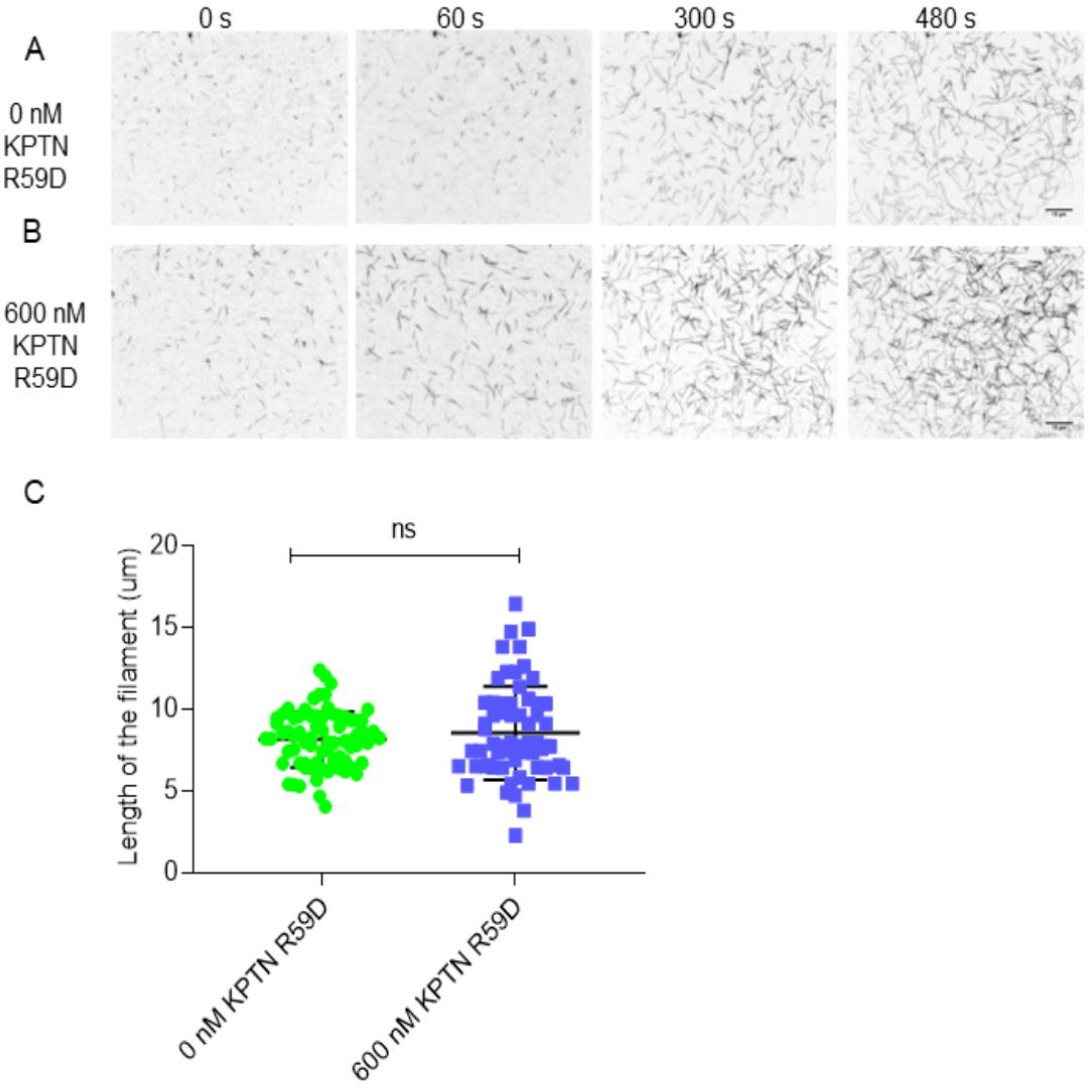
R59D KPTN mutant is incompetent to inhibit actin filament capping. Time-lapse TIRF microscopy images of seeded actin assembly with 333 nM actin seeds and 400 nM actin monomers (barbed end kinetics) in the presence of (A) 0 nM hKPTN^(R59D)^ and (B) 600 nM hKPTN^(R59D)^. Imaging was started after 40 seconds of the reaction mixture in the flow chamber. Scale Bar 10 µm. (C) No significant difference was observed in the length of the actin filaments in 0 nM and 600 nM of _hKPTN(R59D)._

Altogether our data highlights the importance of (Arg^59^) residue in hKPTN to modulate the *in vitro* actin cytoskeleton dynamics.

## Discussion

Multiple actin-binding proteins at the cellular leading edge play a crucial role in regulating actin dynamics, either facilitating or halting filament growth to maintain cellular function and homeostasis. In this study, we identify KPTN, a protein associated with intellectual disability and deafness, as a novel cytoskeleton regulator modulating the actin dynamics. Our research demonstrates that hKPTN possesses barbed end-capping activity. Furthermore, we characterized hKPTN as a member of the WD-repeat-containing protein family, similar to coronin. We also identified its actin-binding site, and its ability to inhibit filament depolymerization. Additionally, hKPTN exhibits weak actin filament bundling activity.

Several earlier reports have discovered KPTN as an actin-associated protein, identified by the monoclonal antibody 2E4, and demonstrated its localization at sites of actin polymerization in various cell types (Bearer 1992a, 1992c, 1993, 1995; Bearer and Abraham, 1999). KPTN was found to be located at the edges of lamellipodia, filopodia, microspikes, and at the tips of the hair cell stereocilia (Bearer et al., 2002). Given KPTN’s peripheral cell localization, it was purified from the cultured fibroblasts and platelet cells *in vitro* using F-actin affinity columns in the presence of ATP (Bearer 1992a).

Previous findings in *Xenopus* have confirmed the upregulation of KPTN expression in embryonic mesoderm prior to early gastrulation movements, implying its crucial role in cell motility (Bearer 1992c). In recent years, several lines of evidence have suggested a role for KPTN in neurodevelopment. Various mutations associated with the KPTN gene have been linked to neurodevelopmental delays, intellectual disability, seizure, macrocephaly, and epilepsy (Baple et al., 2014; Pajusalu et al., 2015; Thiffault et al., 2018; Miguez et al., 2020; Horn et al., 2023; Ullah et al., 2022; Levitin et al., 2023). Recently, the KICSTOR complex, composed of KPTN, ITFG2, C12orf66, and SZT2 was unveiled, which regulates the mTORC1 pathway (Wolfson et al., 2017). A new study has demonstrated a direct link between aberrant mTOR signaling and KPTN mutations, establishing the role of KPTN in the mTOR pathway (Levitin et al., 2023). However, the detailed mechanistic function of KPTN within cells remains largely unknown.

We demonstrated *in vitro-*reconstituted hKPTN binds to actin filaments. Through TIRF microscopy and fluorometric actin nucleation assays, we revealed that hKPTN inhibits actin filament growth in a concentration-dependent manner. We determined the affinity of hKPTN for actin filaments by measuring the length of actin filaments over time. Our study uncovered the barbed end-capping activity of hKPTN, confirmed by biochemical assays with purified hKPTN and actin. Unlike the well-characterized barbed end-capping protein CapZ, our investigations showed that hKPTN functions as a weak barbed end-capping protein. Our analysis suggests that hKPTN prevents filament depolymerization but is not as potent an inhibitor as CapZ. In the presence of Profilin, which supplies ATP-actin monomers to the barbed end (Pernier et al., 2016), our data indicates actin filament elongation, albeit to a limited extent, in the presence of hKPTN. This activity of hKPTN aligns with the fact that KPTN elutes from the F-actin affinity column in the presence of ATP (Bearer 1992a). Other barbed end capping proteins such as Fhod1, Delphilin, and Cdc12p which contain proline-rich FH1 domains allow actin filament elongation in the presence of Profilin (Schönichen et al., 2013; Dutta et al., 2017; Kovar et al., 2003). The mechanisms of filament elongation by formins (those that act as barbed end-capping proteins) and hKPTN in the presence of profilin might differ. In the future, focused studies may provide detailed insights into possible mechanistic differences between these protein activities.

Fluorescently labeled hKPTN was observed at the tip of the actin filament, restricting its growth and implicating its capping activity. hKPTN tended to form thin actin bundles with actin filaments which was not observed in the case of actin-only control reactions. In summary, we characterize hKPTN as an actin filament-binding protein with the capping and bundling activity. The arrangement of actin bundles largely depends upon the architecture of the bundling protein (Winder et al., 2005). The weak actin-bundling activity of hKPTN could be attributed to the oligomeric structure of hKPTN which contains a single binding domain in each subunit or possibly due to the presence of multiple actin-binding domains within its sequence. To gain a deeper understanding of hKPTN’s bundling activity, detailed studies on hKPTN are required.

To investigate its actin-binding site, we conducted a thorough analysis of the hKPTN protein sequence and its predicted AlphaFold structure (Jumper et al., 2021; Tunyasuvunakool et al., 2021). Our data identify KPTN as a member of the WD-repeat-containing protein family closely related to Coronin (Appleton et al.,2006). For the first time, we delineated the actin-binding site in the KPTN protein, which is structurally conserved across various higher eukaryotes. Recent literature has also demonstrated that KPTN, as a member of the KICSTOR complex, is present in all higher eukaryotes (Wolfson et al., 2017). The (Arg^59^) in hKPTN is the crucial amino acid residue that is conserved and critical in regulating actin dynamics. Subsequently, a detailed characterization of the domains within KPTN could be attempted to understand its comprehensive function inside the cells.

The presence of KPTN at the cell periphery and leading edge of the migrating cells is well-documented (Bearer et al., 2002). Our biochemical findings establish the key regulatory activities of hKPTN as a barbed end-capping protein that forms thin actin bundles (as shown in Fig 9). These activities are expected to be central to the reorganization of the actin cytoskeleton at the cell boundary.

**Figure 9:**
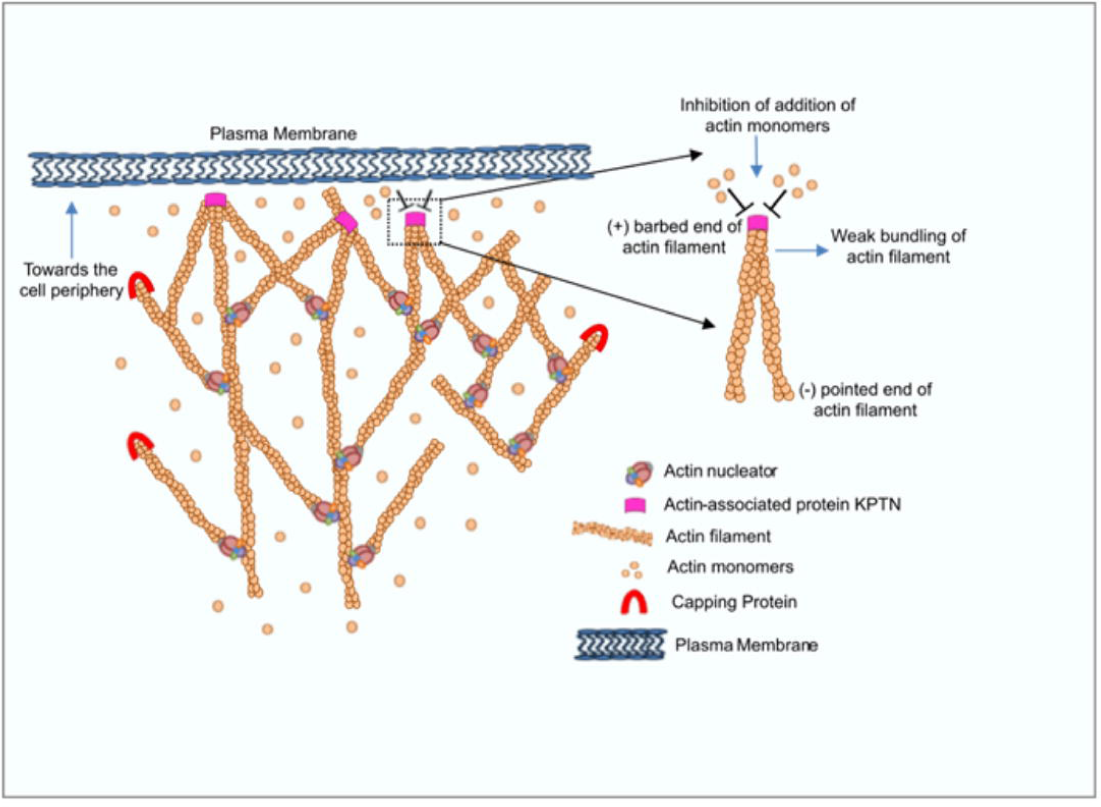
Schematic model figure representing hKPTN localization inside the cell with the actin filaments. The model exhibits the presence of hKPTN towards the periphery of the cell. KPTN binds to the barbed end of the actin filaments and inhibits the barbed end elongation of actin filaments. hKPTN forms thin actin bundles inside the cell. The actin network inside the cell is maintained by the actin nucleator and capping protein.

Actin assembly and disassembly play a vital role in regulating the actin dynamics at the outer edge of the cell. These activities drive critical developmental and homeostatic processes and are tightly regulated by various actin-binding proteins. This study is a significant contribution that describes the mechanistic function of KPTN, a protein well-known for its localization with actin at the cell periphery and its association with neurodevelopmental disorders. Based on these results, future studies on KPTN along with its interactors and other actin regulators are important to comprehensively understand its molecular function as a novel cytoskeleton modulator.

## Material and Method

### Plasmid Construction

The human KPTN (hKPTN) plasmid was purchased from Addgene (Catalogue no.: 100511). The KPTN construct (1-420) aa was subcloned in the pET28a vector (Novagen) using the following primer sequence (Table 1) having BamHI and SalI overhangs. The same construct was cloned in the pSNAP vector (NEB N9181S). The other fragments of KPTN (1-258) aa and (301-420) aa was subcloned in both pET28a and pSNAP vector. The mouse profilin construct was cloned in the pGEX-4T3 vector. The mouse capping protein α subunit and β subunit were cloned separately in the pET28a vector (Dutta P et al., 2017). The Xenopus cofilin in pGEX (GST-Xac1) vector was a kind gift from Prof JR Bamburg (Abe et al.,1996).

### Site-Directed Mutagenesis

The (1-420) aa clone was used as the template with a specifically designed forward and reverse primer (Table 1). The PCR was done using the Phusion polymerase (NEB). The non-mutated amplicons were digested using DpnI (NEB). The mutation was further confirmed by sequence analysis.

### Protein Purification

G-actin was purified from rabbit muscle acetone powder. Actin was labeled with N-(1-pyrene iodoacetamide) [P-29 Molecular probes] and Alexa-488 maleimide [Invitrogen] following the standard protocol (Pollard 1984; Higgs and Pollard 1999; Hansen et al., 2013; Dutta et al., 2017). The labeled and unlabelled actin was purified using an S-200 16/60 gel filtration column pre-equilibrated with G-buffer (2 mM Tris pH 8.0, 0.2 mM DTT, 0.2 mM CaCl_2,_ 0.2 mM ATP) (Hansen et al., 2013). The labeling percentage of the actin was calculated following the standard protocol (Hansen SD et al., 2013) (Juanes et al., 2020).

The 6X-His Tag KPTN and SNAP-KPTN-6X His tag constructs were transformed in BL21 C43(DE3) cells. The protein expression was induced by 0.4 mM IPTG when the O.D. reached 0.6. Subsequently, the cultures were grown for 12 hours at 19°C and harvested at 4000 rpm. The pellets were stored in a −80 freezer. All the steps of protein purification were done on ice following standard protocol. The cells were resuspended in a lysis buffer (50 mM Tris pH 7.5, 150 mM NaCl, 30 mM Imidazole, 1 mM DTT, 0.5% IGEPAL and 1X protease inhibitor cocktail [PIC]) (Dutta et al., 2019). The cells were lysed by sonication and cleared by centrifuging at 14000 rpm for 30 mins. The clear supernatant was incubated with a 50% slurry of Ni-NTA agarose beads (Qiagen) for 2 hours at 4°C. After this, the beads were washed with wash buffer (50 mM Tris pH 7.5, 300 mM NaCl, 30 mM Imidazole). Protein was eluted from the beads using elution buffer (50 mM Tris pH 7.5, 100 mM NaCl, 350 mM Imidazole, and 5% Glycerol). Further, the protein was purified using Superose 6 10/300 Increase column (GE Healthcare) equilibrated with 20 mM Tris pH 7.5, 150 mM NaCl, and 0.5 mM EDTA. For the SNAP-tag KPTN construct, the above protocol remains the same. In the case of the SNAP-tag construct, Ni-NTA beads incubation was done overnight and 20 µM SNAP surface 549/488 (NEB) was added. The excess free dye was removed during the washing of the beads (Breitsprecher et al., 2012).

Mouse Profilin construct was transformed into BL21 DE3 cells. The cells were grown to reach O.D. 0.6. The protein expression was induced by 0.4 mM IPTG and kept at 37°C for 4 hours. The cells were harvested at 4000 rpm and stored at −80°C freezer. The same method was followed to prepare the pellets of mouse capping protein and Xenopus Cofilin. All the steps of protein purification were done on ice following the standard protocol (Dutta et al., 2017). The profilin and Xenopus cofilin pellets were resuspended in lysis buffer (50 mM Tris pH 8.0, 150 mM NaCl, 1 mM DTT, 1 mM EDTA, 1X PIC, and 0.5% IGEPAL), and lysed by sonication then cleared by centrifugation at 14000 rpm for 30 mins at 4°C. The cleared supernatant was incubated with the 50% slurry of Glutathione Agarose (Pierce Thermo) for 2 hours at 4°C. After 2 hours the beads were washed with wash buffer (50 mM Tris pH 8.0, 300 mM NaCl, 1 mM DTT, and 1 mM EDTA). The protein was eluted from the beads using elution buffer (50 mM Tris pH 8.0, 5% Glycerol, and 10 mM Glutathione). The pellets of mouse capping protein were resuspended in a lysis buffer (50 mM Tris pH 8.0, 150 mM NaCl, 30 mM Imidazole, 1 mM DTT, 1X PIC, and 0.5% IGEPAL). The α and β subunit pellets were resuspended in lysis buffer and mixed prior to sonication. The cleared supernatant after sonication and centrifugation was incubated with the 50% slurry of Ni-NTA Agarose beads (Qiagen) for 2 hours at 4°C. The beads were washed with wash buffer (50 mM Tris pH 8.0, 150 mM NaCl, 30 mM Imidazole). The protein was eluted from the beads using elution buffer (50 mM Tris pH 8.0, 100 mM NaCl, 350 mM Imidazole, and 5% Glycerol). All the proteins (Profilin, Capping Protein [CapZ], and Xenopus Cofilin were further purified by Superdex 200 10/300 column and aliquots were flash frozen and stored in the −80°C freezer.

The concentration of all the purified proteins was determined by densitometry of Coomassie-stained bands on SDS-PAGE gels compared to BSA standards. The degree of labeling of the SNAP-tag KPTN was measured by measuring fluorophore absorbance in the solution using the extinction coefficients: SNAP surface_549_ 140,300 cm^-1^ M^-1^; and SNAP surface_488_ 73000 cm^-1^ M^-1^. (Breitsprecher et al., 2012; Henty-Ridilla et al., 2016; Juanes et al., 2020).

### Co-sedimentation Assay

Filamentous actin was formed from 30 µM G-actin for 2 hours at 25 °C with the F-buffer (10 mM Tris-Cl pH 7.5, 0.2 mM DTT, 0.7 mM ATP, 50 mM KCl, 2 mM MgCl_2_). 2 µM hKPTN^(WT)^ and hKPTN^(R59D)^ were incubated for 15 minutes with increasing concentration of actin filaments. For every reaction setup, there was a reaction set for the protein control only. The samples were centrifuged at 310×1000g for 30 minutes using the TLA-100 rotor (Beckman Coulter). The samples were analysed on coomassie-stained 10% SDS PAGE (Dutta et al., 2017).

### Bulk Polymerization Assay

In the case of pyrene-labeled actin assembly, 3 µM of gel-filtered actin monomers (10% labeled) in the G-buffer was incubated with MgCl_2_ and EGTA to convert into Mg-ATP-Actin. The Mg-ATP-Actin was mixed with different concentrations of protein or buffer control (HEKG_5_). For the bulk polymerization assay, the protein was desalted using the PD10 column in HEKG_5_ (20 mM HEPES pH 7.5, 1 mM EDTA, 50 mM KCl, and 5% Glycerol). The polymerization was initiated by adding 3 µl of 20X IM Mix (1M KCl, 40 mM MgCl_2_, 10 mM ATP). Fluorescence was recorded with excitation at 365 nm and emission at 407 nm at 25°C using a fluorescence spectrophotometer (QM40, Photon Technology International, Lawrenceville, NJ) (Dutta et al., 2017, Moseley et al., 2006).

### TIRF Microscopy

Glass coverslips (Corning) and glass slides (Himedia) were cleaned by sonication in 1% HCl followed by acetone, 1M KOH, and ethanol wash for 60 mins in each. The coverslips were rigorously washed in water and air-dried using the hot air stream. The perfusion chambers were prepared from glass slides and coverslips using double-sided sticky tape, and the chambers were coated with silane–PEG (Creative Works) following standard protocols (Portran et al., 2013, Kundu et al., 2021). The samples were imaged using an inverted microscope (ApoN/TIRF 100X/1.49 oil immersion objective on an IX 81 system; Olympus) equipped with a Cell TIRF module (Olympus) and a Hamamatsu ORCA-R2 CCD camera in TIRF buffer (0.2% methylcellulose [4000 cP], 3 mg/ml glucose, 20 µg/ml catalase and 100 µg/ml glucose oxidase), 1% bovine serum albumin and HEKG_5_ buffer. The actin monomer and actin filament concentration and the protein concentration used for TIRF microscopy are specified in the figure legend (Kundu et al., 2021, Hoeprich et al., 2021).

### Image Analysis, Graphical Representation, and Statistical Analysis

Images were processed in FIJI version 1.8.0_322/1.53t (Schindelin et al., 2012) with a 50-pixel rolling-ball radius background subtraction. Actin assembly experiments (nucleation, elongation, and depolymerization kinetics) were visualized in 10 s intervals and dynamic parameters (i.e., elongation rates) were determined as demonstrated in (Henty-Ridilla, 2022). The growth of the actin filament was calculated from traces of actin filament length versus time.

The graphical representation and statistical analyses were performed using GraphPad Prism 8. In the text, we have mentioned the values as mean ± SEM. Graphical representations are scatter dot plots, with the longer middle line indicating the mean and error bars indicating the SEM. The column graphs represent means, and the error bars represent the SEM. unless otherwise indicated. The number of data points calculated in each graph was explained either in the relevant text or the figure legends. The sample sizes were based on the typical number of replicates used in similar studies. An unpaired t-test and a one-way non-parametric ANOVA test were performed to analyze the TIRF microscopy data we obtained for the hKPTN.

## Supporting information

Supplementary figures with leagends

## Availability of Data and Materials

All data generated or analyzed during this study are included in this published article. The raw data are available from the corresponding author on reasonable request.

## Authors Contributions

Conceptualization: PD, AG, and SM. Methodology: PD and IM. Data analysis: PD, IM, and SM. Writing of manuscript: SM, PD, AG and RC. Reagents and Resources: PD, AG, RC, and SM. Project Administration: SM. Funding Acquisition: SM, AG, RC, and PD. All authors gave final approval for publication and agree to be held accountable for the work performed therein.

## ACKNOWLEDGMENTS

The authors are grateful to the facilities of IISER Kolkata, IISER Pune, and NCCS Pune. The authors acknowledge the IISER Pune Microscopy Facility and the National Facility for Gene Function in Health and Disease (NFGFHD) at IISER Pune for access to the equipment and infrastructure. We thank Dr. Srikanth Rapole of the National Centre for Cell Science (NCCS) for the Mass Spectrometry Facility with MS analysis. We acknowledge the help of Shubham Das in plotting the K_d_ calculation graph. The authors thank Dr. Bidisha Sinha and Tanmoy Ghosh of IISER Kolkata for the preliminary TIRF Microscopy experiments.

