## Supplementary figures with leagends for "The actin-associated protein Kaptin modulates F-actin barbed-end dynamics"

**Table 1**

| Primer Name | Sequence |
|---|---|
| KPTN 1st aa Fwd | 5'GGAGCGGATCCATGATGGGGGAGGCGGC3' |
| KPTN 420th aa Rev | 5'GGCACGCGTCGACTTACTGTAGCCGACGTCTCCTCTG3' |
| KPTN R59D Fwd | 5'GGCTTCCGCTACCAAGACCTCGACCAGAAAATCCGGCCAGTG3' |
| KPTN R59D Rev | 5'CACTGGCCGGATTTTCTGGTCGAGGTCTTGGTAGCGGAAGCC3' |

#### Supplementary Figure and Legends:

Supplementary Figure 1

A

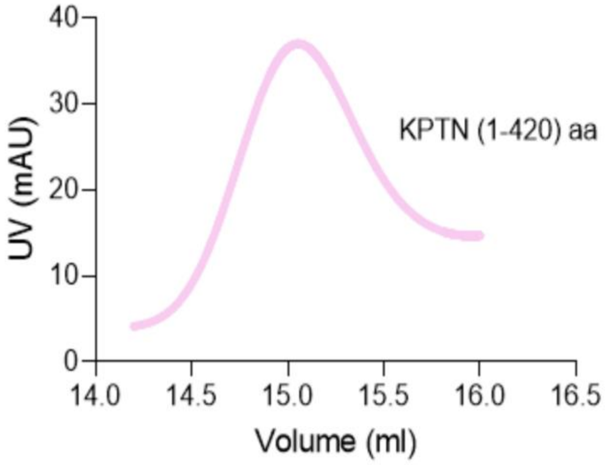


B

MMGEAAVAAGPCPLREDSFTRFSSQSNVYGLAGGAGGRGELLAATLKGKVLGFRYQDLRQKIRPVAKELQFNYIPVDAEI
VSIDTFNKSPPKRGLVVGITFIKDSGDKGSPFLNIYCDYEPGSEYNLDSIAQSCLNLELQFTPFQLCHAEVQVGDQLETV
FLLSGNDPAIHLYKENEGLHQFEEQPVENLFPELTNLTSSVLWLDVHNFPGTSRRLSALGCQSGYVRRVAHVDQRSREVL
QMWSVLQDGPISRVIVFSLSAAKETKDRPLQDEYSVLVASMLEPAVVYRDLLNRGLEDQLLLPGSDQFDSVLCSLVTDVD
LDGRPEVLVATYGQELLCYKYRGPESGLPEAQHGFHLLWQRSFSSPLLAMAHVDLTGDGLQELAVVSLKGVHILQHSLIQ
ASELVLTRLRHQVEQRRRRLQGLEDGAGAGPAENAAS

C

| Protein FDR Confidence | Description | Coverage [%] | Peptides | Unique Peptides | Score Mascot |
|---|---|---|---|---|---|
| High | KICSTOR complex protein kaptin OS=Homo sapiens OX=9606 GN=KPTN PE=1 SV=2 | 65 | 28 | 20 | 6320 |
| High | Keratin, type II cytoskeletal 78 OS=Homo sapiens OX=9606 GN=KRT78 PE=1 SV=2 | 11 | 6 | 4 | 165 |
| High | Keratin, type II cytoskeletal 1b OS=Homo sapiens OX=9606 GN=KRT77 PE=1 SV=3 | 15 | 9 | 5 | 459 |
| High | Albumin OS=Homo sapiens OX=9606 GN=ALB PE=1 SV=2 | 13 | 7 | 7 | 308 |
| High | Desmoglein-1 OS=Homo sapiens OX=9606 GN=DSG1 PE=1 SV=2 | 11 | 7 | 7 | 283 |

Figure S1: **In-gel digestion and Mass spectrometry analysis of hKPTN.** (A) The size exclusion chromatography of hKPTN to get single band purified protein. (B) The arrows mark the start and end residues of the cloned and expressed KPTN. The yellow color-highlighted region represents the overlapping fragments obtained as hits in Orbitrap MS analysis. The unlabelled region represents the region which has not been covered in mass spectrometry. (C) The query coverage map shows 65% matching of the in-gel digested peptides of KPTN protein.

### Supplementary Figure 2

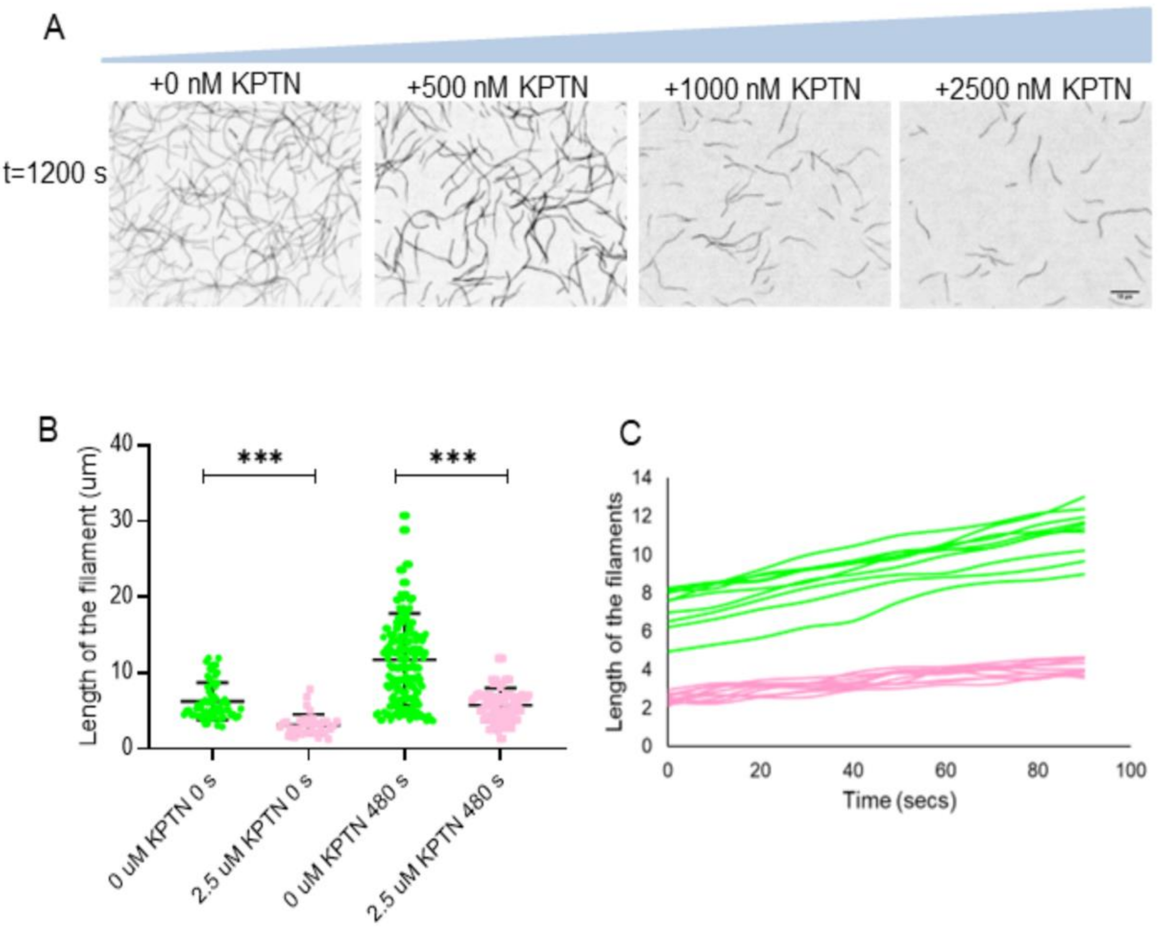

Figure S2: h**KPTN forms thin actin bundles.** (A) Time-lapse TIRF microscopy images of 3000 nM actin monomers. 3000 nM actin monomers were imaged for 480 seconds in the presence of (B) 500 nM KPTN and (C) 2500 nM KPTN. (D) Actin filament bundling skewness was measured at 480 seconds from TIRF images captured with 3000 nM actin monomers as in A, B, and C for five independent times.

Supplementary Figure 3

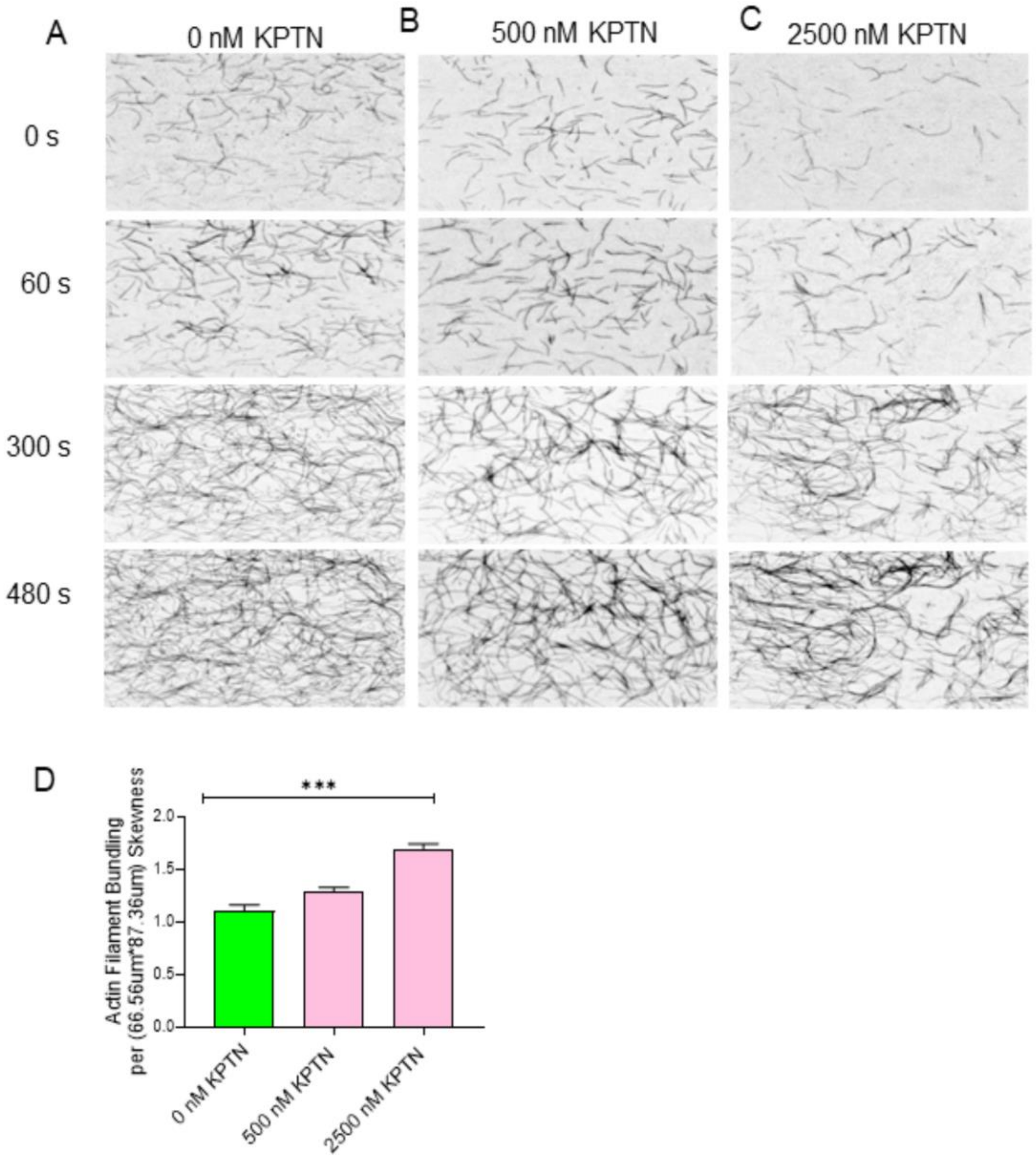

Figure S3: **hKPTN inhibits actin assembly even after 1200 seconds of the reaction.** (A) TIRF microscopy images of 1000 nM actin monomers with increasing concentration of KPTN after 1200 seconds. (B) The average length of the actin filaments was measured at 0 seconds and at 480 seconds for 0 nM and 2500 nM hKPTN. (C) Length of the actin filaments over time plotted for actin control (green) and KPTN (pink) up to 100 seconds. The experiment was repeated thrice.

Supplementary Figure 4

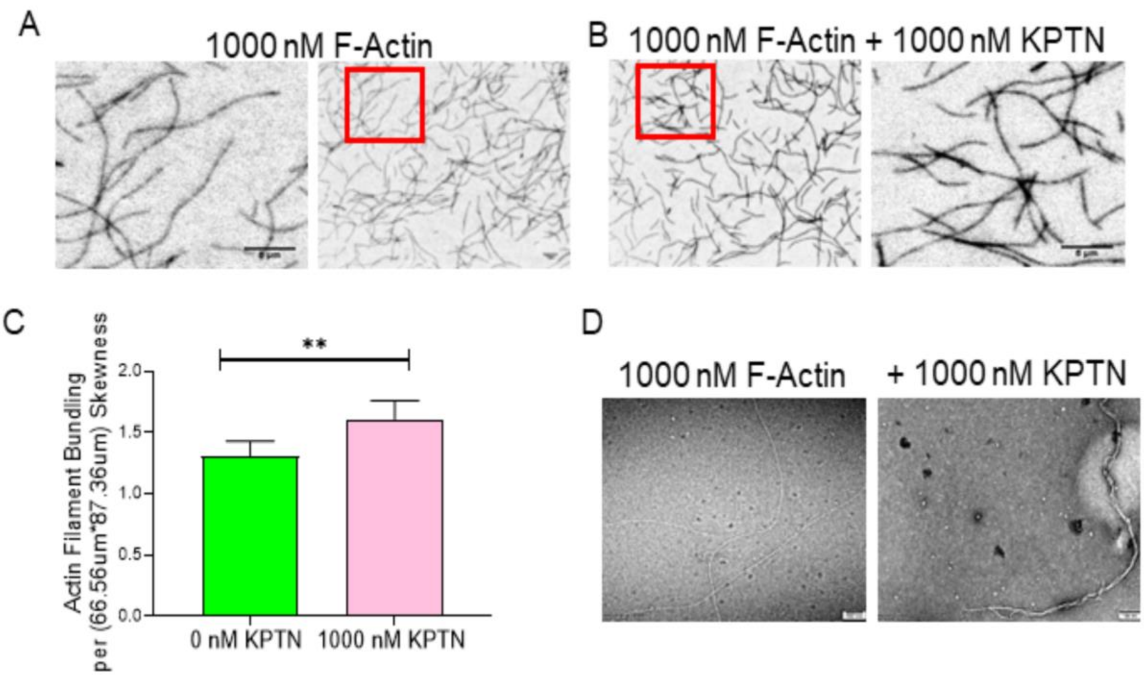

Figure S4: **High resolution of thin actin bundles.** (A) Rhodamine phalloidin stabilized actin filaments were imaged with buffer control. The inset was represented as a zoomed image on the left of the main image. (B) Rhodamine Phalloidin stabilized actin filaments were imaged with 1000 nM of hKPTN. The inset was represented as a zoomed image on the right side of the main image. (C) Actin filament bundling skewness was measured for actin filaments control and actin filaments plus hKPTN. In each case, 10 different fields of view from three independent experiments were considered. (D) High-resolution TEM images showed the thin actin bundles in the case of KPTN but not present in actin control. TEM imaging was performed twice.

Supplementary Figure 5

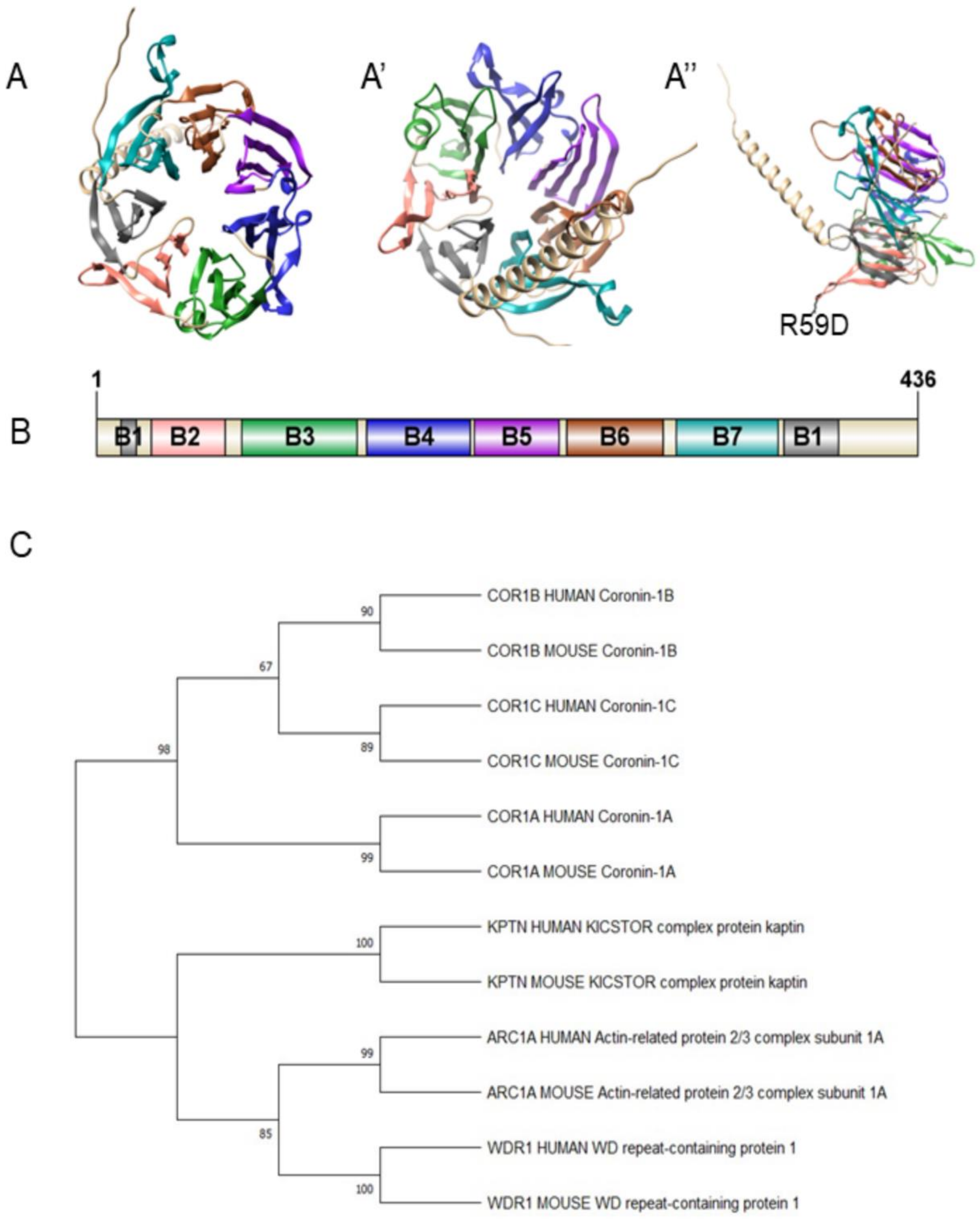

Figure S5: **Predicted 7-bladed beta-propeller structure of KPTN from Alpha fold.** (A) The top-view of the alpha fold predicted beta-propeller structure of hKPTN showed 7 blades containing beta sheets. (A’) The bottom view of the alpha fold predicted the structure of hKPTN. (A’’) The position of R59 amino acid which binds to actin filaments. (B) The schematic representation of the 7-bladed beta-propeller structure of hKPTN. (C) The phylogenetic analysis of beta-propeller structured proteins which bind to actin along with 7-bladed beta-propeller structure of KPTN. The phylogenetic analysis showed the KPTN to be a nearby member of Coronin-1. Only higher eukaryotes were considered for doing the phylogeny as KPTN being a part of KICSTOR is not present in the lower eukaryotes.

### Supplementary Figure 6

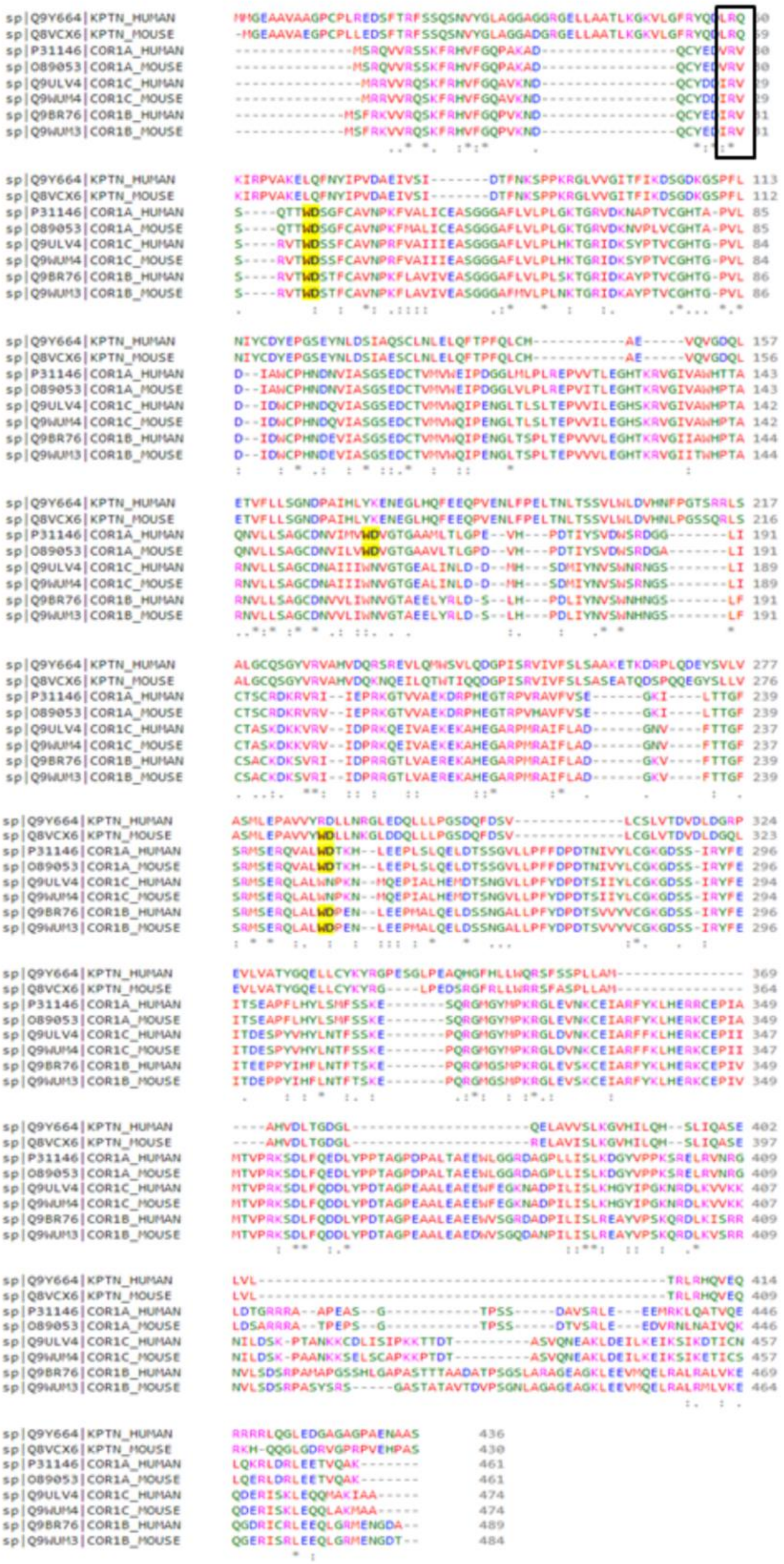

Figure S6: **KPTN is a WD-repeat-containing protein.** The sequence analysis of KPTN with Coronin-1. The R28, R29, and R30 position of Coronin is responsible for actin-binding. This “R” is conserved in the case of KPTN. For hKPTN the “R” position is at 59 (Highlighted in black box). The yellow colour highlighted amino acid showed the presence of WD repeats in Coronin and KPTN.

### Supplementary Figure 7

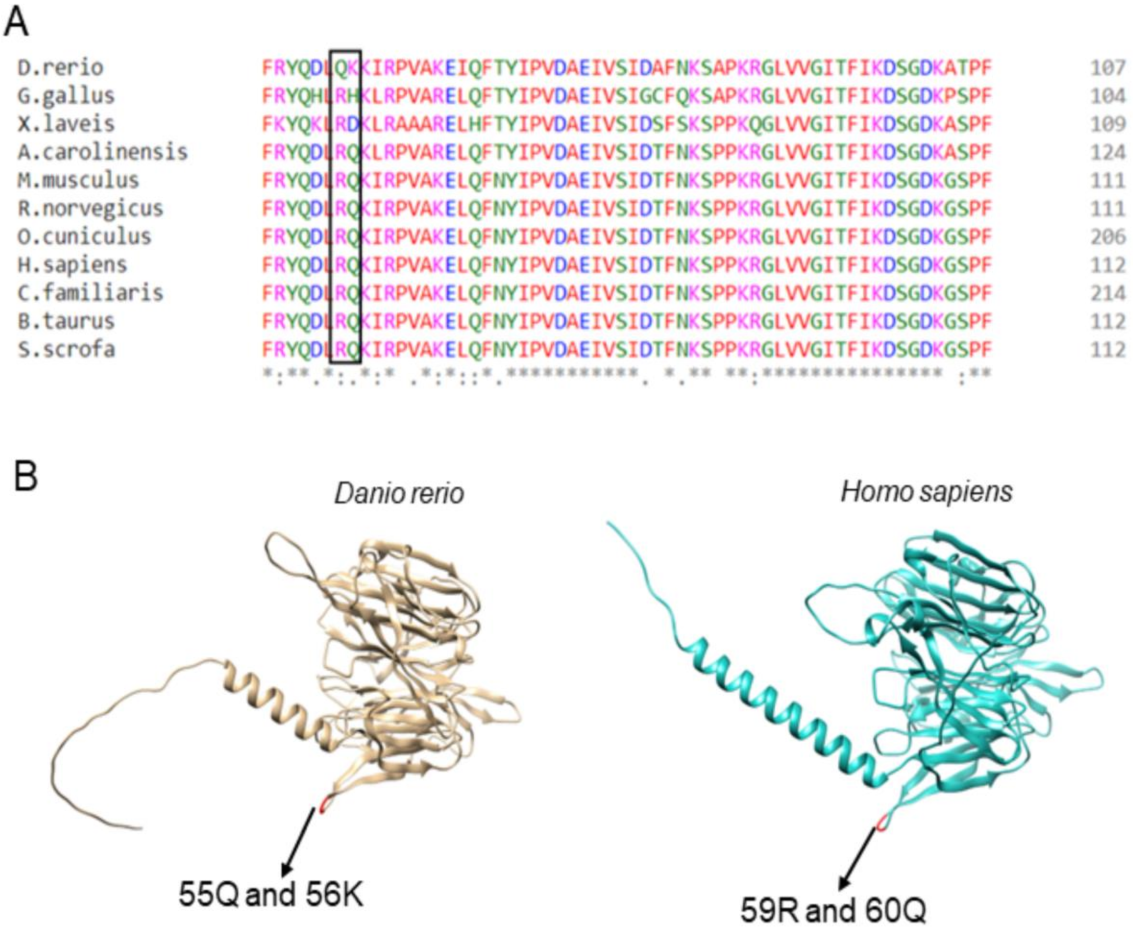

Figure S7: **R59 amino acid is conserved in higher eukaryotes.** (A) The positively arginine amino acid (which is responsible for actin-binding) followed by neutral amino acid is structurally conserved in the higher organisms. (B) In the case of *Danio rerio,* the positively charged amino acid and neutral amino acid interchanged their position but they are structurally conserved with other higher organisms.

**Movie Legends:**

Supplementary Movie 1**: (Associated with Fig 2A) Assembly of Actin Filaments.** The reaction mixture contains 1000 nM actin monomers (15% Alexa-488 labeled). No KPTN was used in the reaction. Scale bar 10 µm.

Supplementary Movie 2**: (Associated with Fig 2B) Actin assembly is inhibited by hKPTN.** The reaction mix had 1000 nM actin monomers (15% Alexa-488 labeled). The concentration of hKPTN in the reaction mix was 500 nM. Scale bar 10 µm.

Supplementary Movie 3**: (Associated with Fig 2C) hKPTN inhibits actin assembly.** The reaction mix had 1000 nM actin monomers (15% Alexa-488 labeled). The concentration of hKPTN in the reaction mix was 1000 nM. Scale bar 10 µm.

Supplementary Movie 4**: (Associated with Fig 2D) hKPTN-mediated hindered actin assembly.** The reaction mix had 1000 nM actin monomers (15% Alexa-488 labeled). The concentration of hKPTN in the reaction mix was 2500 nM. Scale bar 10 µm.

Supplementary Movie 5**: (Associated with Fig 2F) hKPTN-mediated actin assembly.** The reaction mix had 1000 nM actin monomers (15% Alexa-488 labeled). The concentration of hKPTN in the reaction mix was 750 nM. Scale bar 10 µm.

Supplementary Movie 6**: (Associated with Fig 2F) Actin assembly by hKPTN.** The reaction mix had 1000 nM actin monomers (15% Alexa-488 labeled). 1500 nM of hKPTN was added to the reaction mix. Scale bar 10 µm.

Supplementary Movie 7**: (Associated with Fig 3A) Actin filament elongation.** The reaction had 400 nM actin monomers (15% Alexa-488 labeled) and 333 nM of unlabeled actin filament seed. Scale bar 10 µm.

Supplementary Movie 8**: (Associated with Fig 3B) Actin filament elongation in the presence of CapZ.** The reaction had 400 nM actin monomers (15% Alexa-488 labeled), 333 nM of unlabeled actin filament seed, and 100 nM of CapZ. Scale bar 10 µm.

Supplementary Movie 9**: (Associated with Fig 3C) Actin filament elongation in the presence of hKPTN.** The reaction had 400 nM actin monomers (15% Alexa-488 labeled), 333 nM of unlabeled actin filament seed, and 300 nM of hKPTN. Scale bar 10 µm.

Supplementary Movie 10**: (Associated with Fig 3D) Actin filament elongation impeded by the presence of hKPTN.** The reaction had 400 nM actin monomers (15% Alexa-488 labeled), 333 nM of unlabeled actin filament seed, and 600 nM of hKPTN. Scale bar 10 µm.

Supplementary Movie 11**: (Associated with Fig 4A) Actin filament elongation in the presence of Profilin.** The reaction had 400 nM actin monomers (15% Alexa-488 labeled), 333 nM of unlabeled actin filament seed, and 3000 nM of Profilin. Scale bar 10 µm.

Supplementary Movie 12**: (Associated with Fig 4B) Actin filament elongation in the presence of Profilin and hKPTN.** The reaction had 400 nM actin monomers

(15% Alexa-488 labeled), 333 nM of unlabeled actin filament seed, 3000 nM of Profilin, and 600 nM of hKPTN. Scale bar 10 µm.

Supplementary Movie 13A**: (Associated with Fig 6B) The actin monomers polymerize into filament.** 1000 nM actin monomers (15% Alexa-488 labeled) were used in the reaction mixture. The colour of actin filaments was changed to red to keep it the same as in Figure 6A. Scale Bar 5 µm.

Supplementary Movie 13B**: (Associated with Fig 6B) The SNAP labeled hKPTN inhibits the actin filament polymerization.** The hKPTN was labeled with SNAP surface 549 (NEB). 1000 nM actin monomers (15% Alexa-488 labeled) were used in the reaction mixture. 2500 nM of hKPTN (30% SNAP Surface 549 labeled) was added in the reaction to observe hKPTN-mediated actin polymerization. The colour of actin filaments and hKPTN were changed to red and green respectively to keep it the same as in Figure 6A. Scale Bar 5 µm.

Supplementary Movie 14A**: (Associated with Fig 7E) Actin assembly by hKPTN$^{(WT)}$.** The reaction was set up with 1000 nM actin monomers (15% Alexa-488 labeled). The flow chamber contains 2500 nM of KPTN$^{(WT)}$. Scale bar 10 µm.

Supplementary Movie 14B**: (Associated with Fig 7E) Actin assembly by hKTPN$^{(R59D)}$.** The reaction was set up with 1000 nM actin monomers (15% Alexa-488 labeled). The flow chamber contains 2500 nM of KPTN$^{(R59D)}$. Scale bar 10 µm.

Supplementary Movie 15**: (Associated with Fig 8A) Actin filament elongation.** The reaction mix had 400 nM of actin monomers (15%-Alexa 488 labeled). This control reaction was performed to compare the elongation assay reaction with the hKPTN$^{(R59D)}$ mutant in Fig 8B. Scale bar 10 µm.

Supplementary Movie 16**: (Associated with Fig 8B) Elongation of actin filaments by hKPTN$^{(R59D)}$.** The reaction mix had 400 nM of actin monomers (15%-Alexa 488 labeled) and 600 nM of hKPTN$^{(R59D)}$. Scale bar 10 µm.

Supplementary Movie 17**: (Associated with Supplementary Fig 3A) Assembly of actin filaments using 3000 nM of actin monomers.** 3000 nM of actin monomers (15% -Alexa 488 labeled) were used in the reaction mixture. Scale bar 10 µm.

Supplementary Movie 18**: (Associated with Supplementary Fig 3B) In the presence of high actin monomer concentration, inhibition of actin assembly by hKPTN.** 3000 nM of actin monomers (15% -Alexa 488 labeled) were used in the reaction mixture. 500 nM of hKPTN was used in the reaction. Scale bar 10 µm.

Supplementary Movie 19**: (Associated with Supplementary Fig 3C) Assembly of 3000 nM actin monomers into filaments in the presence of hKPTN.** 3000 nM of actin monomers (15% -Alexa 488 labeled) were used in the reaction mixture. 2500 nM of hKPTN was used in the reaction. Scale bar 10 µm.
